# Thermal stress disrupts microbial symbiosis and host activity in the model sponge *Ianthella basta*

**DOI:** 10.64898/2026.10.01.756110

**Authors:** J. Pamela Engelberts, Laura Rix, Heidi M. Luter, Bettina Glasl, Nico Jehmlich, Katharina Kitzinger, Ira Cooke, Anton Legin, Craig W. Herbold, Maximilian Marini, Katherine A. Dougan, Steven J. Robbins, Michael Wagner, Nicole S. Webster

**Affiliations:** Centre for Microbiome Research, School of Biomedical Sciences, Queensland University of Technology (QUT), Translational Research Institute, Woolloongabba, QLD, Australia; Australian Centre for Ecogenomics, School of Chemistry and Molecular Biosciences, The University of Queensland, Brisbane, QLD, Australia; Australian Institute of Marine Science, Townsville, Queensland, Australia; Centre for Microbiology and Environmental Systems Science, Division of Microbial Ecology, University of Vienna, Vienna, Austria; Department of Molecular Systems Biology, Helmholtz-Centre for Environmental Research - UFZ, Leipzig, Germany; Centre for Tropical Bioinformatics and Molecular Biology, James Cook University, Townsville, Queensland, Australia; School of Biological Sciences, University of Canterbury, Christchurch, New Zealand; Center for Microbial Communities, Department of Chemistry and Bioscience, Aalborg University, Aalborg, Denmark; University of Tasmania, Hobart, TAS 7004 Australia

## Abstract

Climate change is threatening coral reef ecosystems around the world, causing significant declines in reef organisms, including sponges. The health of sponges is intimately linked to their symbiosis with microorganisms, yet little is known about their integrated response to ocean warming. Here, we present a comprehensive view of the effect of elevated sea surface temperatures (SST) on the model sponge *Ianthella basta. I. basta* individuals were exposed to temperatures up to 32 °C, 2-3 °C above the mean annual maxima. Host gene expression profiles showed an initial response to heat stress after 2 days, and a decreased expression of genes involved in maintaining tissue integrity, pumping capacity, vitamin transport, and stable symbioses after 4 to 8 days. Congruently, heat stressed samples started to show tissue regression and a reduced abundance of the dominant symbionts after 4 days. A significant loss of the symbionts occurred after 8 days, together with a reduction in microbially driven nitrogen cycling and amino-acid transport and an introduction of opportunistic microbes, which could lead to nutrient imbalance and further destabilisation of the holobiont. Together, this work provides important insights into the avenues through which future SST can impact host-microbe interactions in coral reef sponges.

## Introduction

Climate change-driven ocean warming is threatening coral reef ecosystems around the world, causing significant declines in essential reef-building organisms [1-4]. With sea surface temperatures (SST) predicted to rise by up to 4 °C within the next century [5], many reef invertebrates will be pushed beyond their current thermal limit, potentially driving coral reefs to extinction [6]. To understand and ultimately predict the ability of reef invertebrates to adapt to increasing SST, research has assessed how they respond to thermal stress [7-10]. However, most of this work has focused on corals and their symbiotic association with dinoflagellates (i.e. Symbiodiniacea), leaving other keystone reef invertebrates, such as sponges, under-studied [11, 12].

Marine sponges are important constituents of coral reefs, where they can occupy up to 30% of the benthic substrate, enhancing the survival of a wide range of species through the provision of microhabitats and settlement surfaces [13, 14]. Moreover, they mediate the retention and recycling of (in)organic nutrients and dissolved organic matter by continuously filtering thousands of litres of seawater [15, 16]. The health of sponges is intimately linked to their symbiosis with microorganisms, which can provide the host with nutrients, and aid in chemical defence and the removal of waste products [17-20]. While shifts in the microbial community have often been linked to a decrease in host health [21-24], the role the microbiome plays in the thermal stress response of sponges (i.e. causative or responsive) remains challenging to ascertain, particularly given the high diversity and complexity of sponge-associated microbial communities. Furthermore, few studies have examined the integrated response of the sponge host and microbiome to increasing SST, hampering the identification of molecular mechanisms that underpin dysbiosis and sponge mortality [21]. Characterising the thermal stress response of each member of the sponge holobiont would increase our understanding of how environmental stress impacts these keystone species and aid in predicting their ability to adapt to increasing SST.

Here, we investigated the effect of elevated seawater temperatures on the model sponge *Ianthella basta. I. basta* is an abundant and widely distributed reef sponge in the Indo-Pacific and hosts only three dominant symbionts: an ammonia-oxidizing-archaeaon (‘*Ca*. Nitrosospongia ianthellae’), a Gammaproteobacterium (‘*Ca*. Taurinisymbium ianthellae’), and an Alphaproteobacterium (‘*Ca*. Luteria ianthellae’) [25-27], allowing for the in-depth characterisation of the response of each holobiont member to thermal stress. Six *I. basta* individuals were exposed to temperatures up to 32 °C, 2-3 °C above the mean annual maxima, and changes in the morphology and activity of the host and in the abundance and metabolism of the microbiome were measured. We show that after 4 days of elevated SST, host tissue started to necrose and symbiotic functions became suppressed, which was followed by a decreased abundance of the dominant symbionts. The breakdown of the holobiont led to the introduction of microbes belonging to taxa which are specialists in the degradation of organic matter. Moreover, the loss of the dominant symbionts resulted in a drastic change in the metabolism of the microbiome, where heat stressed samples showed a reduction in microbially driven nitrogen cycling and amino-acid biosynthesis and transport, as well as a reduced ability to persist in the host tissue, further disrupting the nutritional interdependence between host and symbionts. Taken together, this work provides unprecedented insight into the heat stress response of the sponge holobiont and shows that SST predicted to occur within the next century can cause a dysbiosis in the symbiotic interactions in *I. basta*, ultimately leading to sponge death.

## Materials and Methods

### Sample collection and experimental setup

The tissue of *Ianthella basta* can be designated as either thin or thick (**Figure 1**) [27]. To account for the effect of tissue thickness on the sponge’s response to heat stress, yellow individuals with both thin and thick tissue were used throughout the experiment. Three *I. basta* individuals with a yellow, thin morphology were collected from Davies Reef (S 18°49.354’, E 147°38.253’) in September 2020, using SCUBA at depths between 12-15 m. A further three *I. basta* individuals with a yellow, thick morphology were collected from the same location and depth using SCUBA in February 2021. After collection, sponges were immediately transferred to a 2,500 L mesocosm in the National Sea Simulator (SeaSim) at the Australian Institute of Marine Science (AIMS) in Townsville, Australia. Each individual was fragmented into 24 equally sized clones and left to acclimatise for 24 h. Fragments were then evenly distributed over six 50 L indoor tanks (three control and three treatment tanks). Each tank was initially maintained at 29 °C, corresponding to the ambient water temperature on Davies Reef in February 2021. Tanks were continuously supplied with 0.4 μm filtered seawater (flow rate = 0.8 L min^−1^, 100% water exchange per 1 h) and kept at a 12 h:12 h light/dark regime (peak daytime irradiation = 80 µmol m^-2^ s^-1^). Supplemental feeding was provided every other day for the duration of the experiment (30 mL of microscopic algae, 2000 cells ml^−1^).

**Figure 1.**
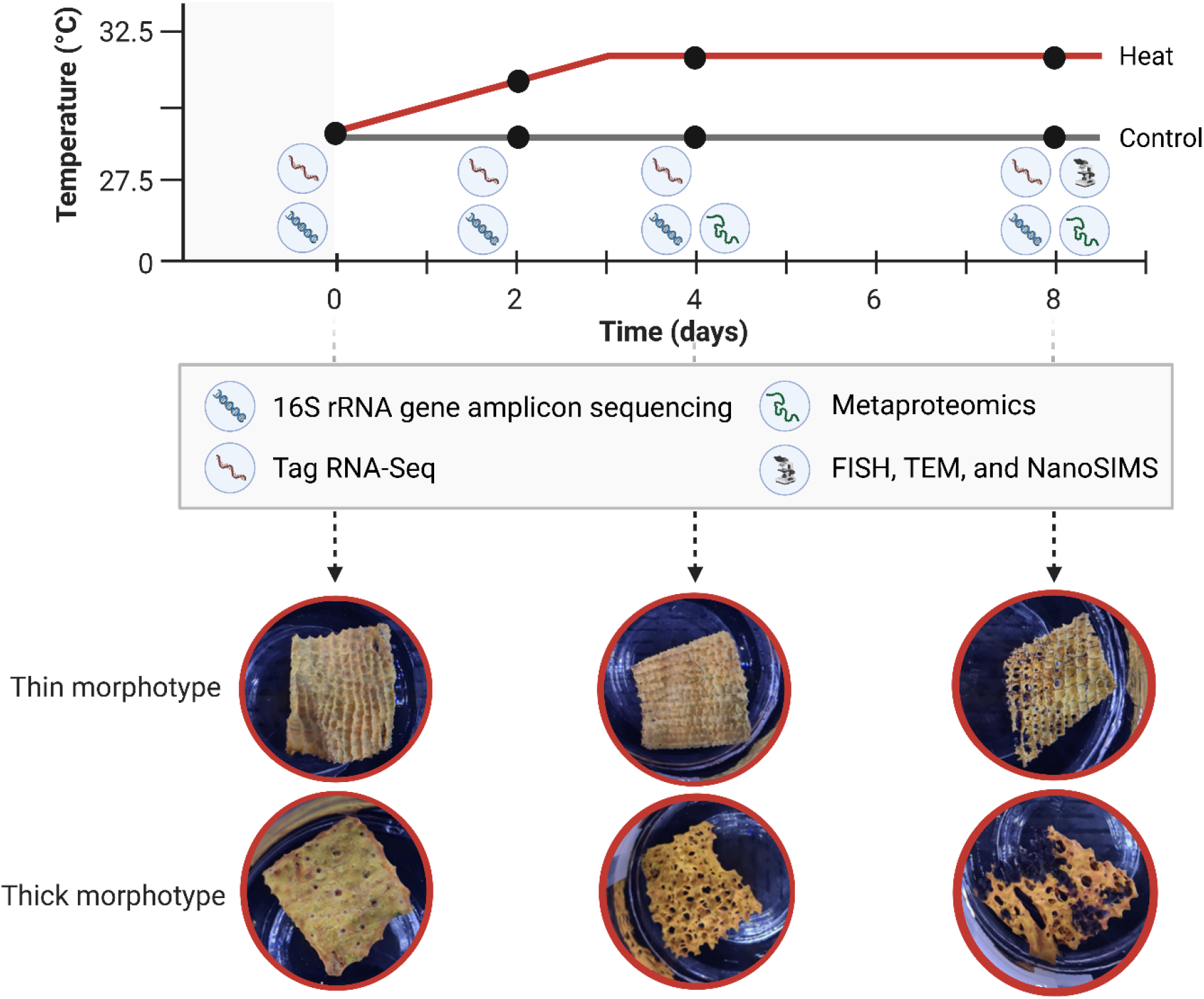
Schematic overview of the *I. basta* morphotypes used in this study, temperature treatments, and sampling time points (represented by the black dots). Photos of *I. basta* clones with a thin (upper) and thick (lower) morphotype show morphological changes observed at timepoint 0, and after 4 and 8 days of heat stress.

After 4 days of acclimation, the temperature in the three treatment tanks was gradually ramped by 0.5 °C per 12 h to a final constant temperature of 32 °C, a temperature previously linked to sponge dysbiosis and discoloration [28-30]. The three control tanks were kept at 29 °C throughout the experiment. One clone of each individual (*n* = 3 per morphotype) was sampled from a control and treatment tank prior to the temperature ramp-up (T = 0 h), and after two days (T = 2 days, Temperature = 31.5 °C), four days (T = 4 days, Temperature = 32 °C), and 8 days of heat stress (T = 8 days, Temperature = 32 °C, **Figure 1, Table S1)**. Upon collection, tissue of each clone was photographed and snap frozen in liquid nitrogen and stored at -80 °C. This tissue was used to analyse the host gene expression profile through Tag RNA-sequencing and the microbial community through 16S rRNA gene amplicon sequencing and metaproteomics. Tissue from the same clone was further collected for fluorescence *in situ* hybridisation (FISH) and transmission electron microscopy (TEM). For FISH, tissue was fixed in 4% PFA overnight and stored in 1:1 ethanol-phosphate-buffered saline (PBS) at -20 °C. For TEM, small pieces of tissue (∼1 mm^3^) were fixed in 3% glutaraldehyde and 1% paraformaldehyde in buffer containing 0.2M sodium cacodylate, 0.35M sucrose, and 0.1M sodium chloride for 4h at room temperature followed by overnight fixation at 4 °C. Samples were subsequently transferred to buffer (0.2M sodium cacodylate, 0.3M sodium chloride) and stored at 4 °C until further processing.

### Tag-seq: RNA extraction, sequencing, and processing

To allow for downstream Tag RNA-seq analyses, a method that generates a single initial library molecule per transcript complementary to 3′-end sequences [31], a host reference transcriptome was constructed for both morphotypes of *I. basta* (thin and thick, **Table S2**, see Supplementary Methods). To characterise the transcriptomic response of *I. basta* to heat stress, total RNA was extracted from 12 control and 12 treatment samples per morphotype, following the same methods as described for the host reference transcriptome. Total RNA quantity and quality was measured using Qubit and Nanodrop and sent to the University of Texas at Austin Genome Sequencing and Analysis Facility for Tag RNA-seq, following the protocols described in Meyer et al. [31] and Lohman et al. [32]. In short, libraries were prepared using 500 ng of total RNA and 10 cycles of cDNA amplification. Pooled libraries were run on a NovaSeq6000 (single-end, 100-base run) at a depth of 10 million reads per sample. Raw reads were trimmed for adapter sequences and quality filtered using cutadapt v2.9 [33], keeping reads with a minimum length of 25 bp and quality score of 15. Reads were subsequently mapped against the corresponding host reference transcriptome (i.e. thin against thin and thick against thick) using bowtie2 v2.4.4 [34] with the -very-sensitive flag. The resulting read counts were assembled per isogroup (∼gene) and morphotype (**Table S3**).

### Differential gene abundance analysis, weighted gene co-expression network analysis (WGCNA), and gene ontology (GO) enrichment analysis

Read count data was normalised using DESeq2 v1.34.0 [35] in R v4.2.1 [36], excluding genes with less than 10 counts over the 24 samples per morphotype from further analyses. The molecular response of the morphotypes to heat stress was subsequently characterised by identifying differentially expressed genes (p < 0.05) between control and treatment samples for each time point using DESeq2 v1.34.0 [35]. To assess the general transcriptomic response with principal component analysis (PCA), counts were further normalised with the variance stabilizing transformation (vst, **Figures S1 and S2**). Due to an observed individual effect (i.e. replicates from the same sponge individual cluster together), the vst transformed data was subsequently used to reconstruct gene co-expression networks using Weighted Gene Co-expression Network Analyses (WGCNA, v1.70-3) [37].

WGCNA is an approach that assigns genes with similar expression patterns to modules. WGCNA can then separate modules that correlate to temperature from modules that correlate to time point and/or individual, allowing for the identification of groups of genes that change expression solely in response to elevated temperature. WGCNA was run for all 24 samples per morphotype after checking for outliers in an unrooted hierarchical tree, which was built with the hclust “average” function in R v4.2.1 [36]. Soft thresholding values of 16 (thin) and 10 (thick) were picked with the WGCNA pickSoftThreshold function to assess gene expression adjacency by reconstructing a topological overlap matrix similarity network. This network was used to identify modules using the WGCNA dynamicTreeCut function, with a minimum module size of 30 genes. Modules that were 70% (thin) and 65% (thick) similar were subsequently merged, which resulted in 17 (thin) and 16 (thick) modules that were used for module-trait correlation and gene ontology (GO) enrichment analysis. Modules were clustered by eigengene dissimilarity using the hclust “average” function, after which the correlation between eigengenes and traits (i.e. temperature, time point, and individual) was calculated. Significant correlation values (p < 0.05) were plotted as a heatmap using the labeledHeatmap function of WGCNA for **Figure S3A** and **B**. The expression profiles of the modules that correlated significantly to temperature were further visualised with ggplot2 v3.3.5 for **Figure S3C** and **D**.

To gain insight into functions that were up- or downregulated in response to heat stress, a GO enrichment analysis was conducted on modules that significantly correlated to temperature using topGO [38]. TopGO identifies over-represented GO groups by comparing, for each GO term, the number of annotations assigned to genes within a module to annotations assigned to the rest of the data set. The significance of enriched GO terms was subsequently tested using Fisher’s exact test (algorithm = ‘elim’). Overlapping GO terms (i.e. GO terms significantly up- or downregulated in both morphotypes) were visualised with ggplot2 v3.3.5 for **Figure 2**.

**Figure 2.**
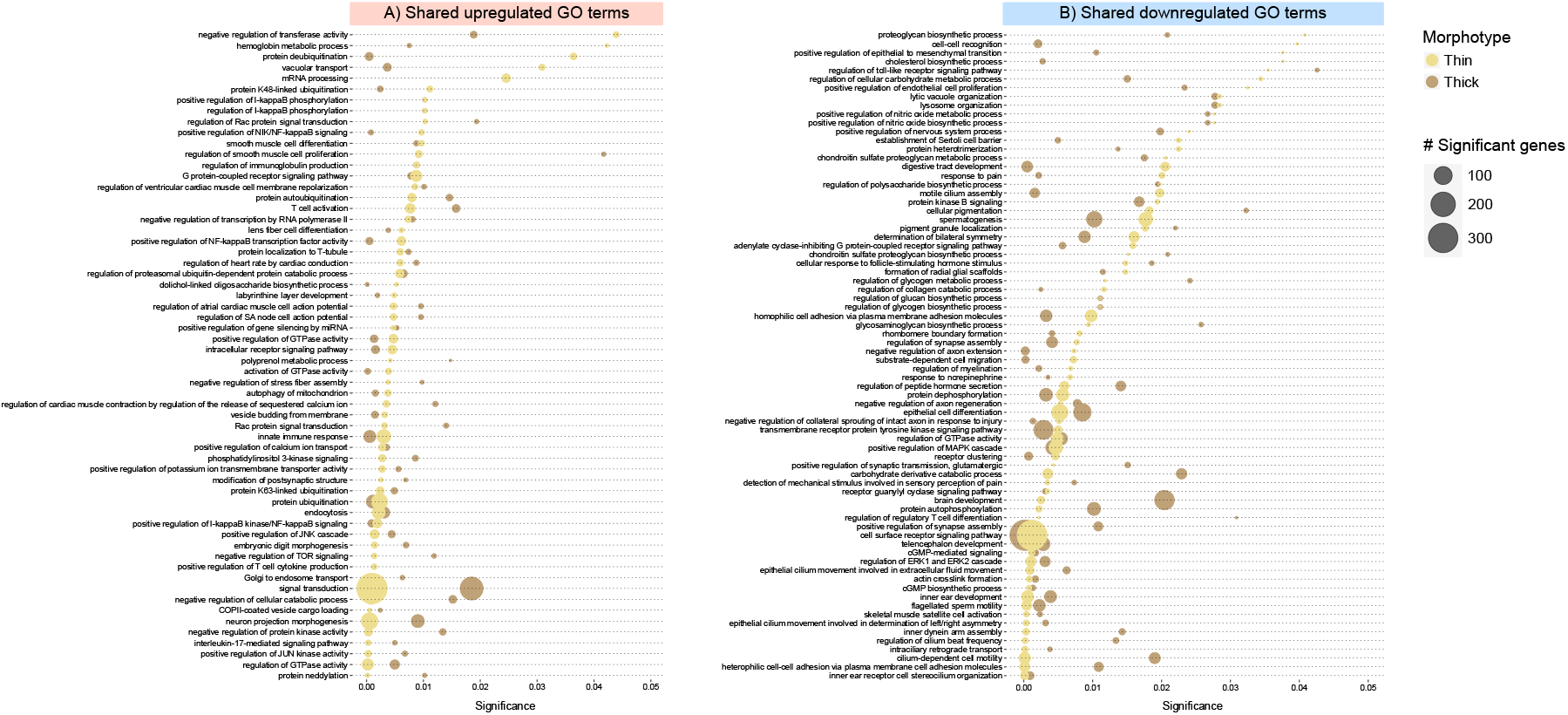
Gene ontology (GO) enrichment analysis showing gene groups that are significantly upregulated (A) and downregulated (B) in both morphotypes in response to heat stress. GO enrichment analysis is based on the Weighted Gene Co-expression Network Analyses (WGCNA) and shows the molecular response of the sponge host to elevated temperature across all time points.

### Microbial DNA extraction, 16S rRNA gene amplicon sequencing, and sequence analysis

DNA was extracted from 48 samples using the DNeasy Powersoil Pro Kit (Qiagen, Germany), following the manufacturer’s instructions, including a bead beating step of 40 s at 2500 rpm. Extracted DNA was sent to the Australian Centre for Ecogenomics (ACE, Brisbane) for sequencing of the V4 region of the 16S rRNA gene, using the primers 515F [39] and 806R [40] targeting both bacteria and archaea. 16S rRNA gene libraries were prepared using the workflow outlined by Illumina (#15044223 Rev.B) followed by sequencing on an Illumina MiSeq Sequencing system (2 × 300 bp).

Raw sequences were quality filtered and trimmed with Trimmomatic [41], keeping sequences with a minimum length of 250 bp and an average base quality above 15. QIIME2 v2020.11.1 [42] was used to process quality controlled reads for feature selection, taxonomic assignment, and abundance calculations. Reads were de-noised (filtered, dereplicated and chimeras identified and removed) using DADA2 (--p-trunc-len = 0) [43] and relative frequencies of each resulting feature calculated. Taxonomy was assigned to features by BLASTing its sequence against the SILVA database (release 138.1, clustered at 99% identity; http://www.arb-silva.de/) using the classify-consensus-blast function with default parameters. Features will be referred to as amplicon sequence variants (ASVs) from here on. Chloroplasts, mitochondrial sequences, and singletons were removed from the dataset, as well as ASVs with an overall abundance below 0.05%.

Statistical analyses were performed in R v4.2.1 [36] and PRIMER 7 [44]. Bray Curtis dissimilarities were calculated for all samples and visualised in a two-dimensional space by Non-metric Multi-dimensional Scaling (NMDS, ‘phyloseq package’, **Figure S5**). To determine whether the microbial community of heat stressed samples was significantly different from control samples, permutational multivariate analyses of variance (PERMANOVAs, 999 permutations) were performed using PRIMER 7. To then identify which ASVs contributed most to the observed differences between treatments, a similarity percentage analysis (SIMPER) was performed on the square root transformed data (to account for rare species).

### Microbial enrichment, protein extraction, and proteome analyses

Samples collected on T = 4 days and T = 8 days were further processed for metaproteomics (6 control and 6 treatment samples per morphotype) to elucidate heat induced changes in the function of the microbiome. To this end, frozen samples were ground on liquid nitrogen and immediately transferred into sterile Calcium/Magnesium-Free Sea Water (CMFSW; 25 g NaCl, 0.8 g KCl, 1 g Na_2_SO_4_, 0.04 g NaHCO3 per 1 L milliQ water) containing Protease Inhibitor Cocktail VI (PIC; 2 µl per ml; A.G. Scientific, US) to preserve cellular protein content. Samples were filtered through a 100 μm Nylon sterile cell strainer and centrifuged for 15 min at 100 × g at 4 °C to remove sponge cells. The supernatant was subsequently centrifuged for 15 min at 300 × g at 4 °C, collected, and filtered through a sterile 8 μm and 5 μm TETP02500 Isopore membrane filter (Merck, Germany). The microbial enriched fraction of the samples was subsequently pelleted with a centrifugation at 8,000 × g for 20 min at 4 °C. The pellet was washed in CMFSW with PIC and centrifuged again at 5,000 × g for 20 min at 4 °C. This last step was repeated twice. The final microbial pellet was recovered in 1 mL TE buffer with PIC and stored at -80 °C until sent to the Centre for Environmental Research, Helmholtz, for metaproteomic analysis. At the Centre for Environmental Research, the microbial cell pellet was dissolved in 1 mL lysis buffer (8M Urea, 2M Thiourea, and 1mM phenylmethanesulfonyl fluoride; PMSF) and proteins were extracted and analysed as described in Engelberts et al. [27].

## Results and Discussion

### Ianthella basta shows tissue regression after 4 days of heat stress

A total of 24 sponge fragments were used throughout the experiment per *I. basta* morphotype (12 control and 12 treatment, **Table S1**). After 4 days of heat stress, all fragments from two thick individuals started to show signs of tissue regression (**Figure 1**), indicating the sponges were stressed. After 8 days of heat stress, all clones displayed tissue regression (thin and thick) and discolouration (thick; **Figure 1**). Control fragments showed no tissue regression over time. The morphological response of the thin morphotype to heat stress was consistent with previous work on a thin *I. basta*, where tissue regression occurred after 7 days at 31 °C and discolouration of the tissue only after 14 days [29]. In contrast, the thick morphotype showed a morphological response almost twice as fast, suggesting that it is less tolerant of increasing SST.

### Recovery of high-quality host reference transcriptomes

To characterise the underlying molecular response of *I. basta* to increasing SST through Tag RNA-seq analyses, two host reference transcriptomes were constructed (one per morphotype). Sequencing of the total RNA from the thin and thick morphotypes yielded 121 and 108 million raw reads, respectively. After trimming, 99 (thin) and 88 (thick) million reads remained, which were assembled into two transcriptomes consisting of 425,800 (thin) and 467,886 (thick) sequences. The transcriptomes had an average sequence length of 675 bp (thin) and 626 bp (thick), and an N50 of 963 (thin) and 854 (thick; **Table S2**). A total of 29,319 (thin) and 38,736 (thick) isogroups (∼genes) were identified in the transcriptomes, of which 97% were functionally annotated (i.e. assigned either a KO or PFAM; **Table S3**). Based on the presence of complete single-copy orthologues of the metazoan BUSCO lineage, the transcriptomes were 93.4% (thin) and 92.9% (thick) complete (**Table S2**), which is higher than most previously published sponge transcriptomes, for which the average completeness is 83.15 ± 12.7% [45-51].

### Transcriptomic response of I. basta to thermal stress: general pattern and initial response

Ordination analysis of the general expression profiles of *I. basta* showed a strong individual effect for the thin morphotype (**Figure S1A**), which was less pronounced in the thick morphotype (**Figure S1B**). After 4 (thick) and 8 (thin and thick) days at 32 °C, samples started to cluster separately from control samples, congruent with the observed morphological response. The thick morphotype further displayed clustering of samples from T = 0h with T = 8 days treatment samples (**Figure S1B**). This clustering can be explained by the fact that clones with a thick morphotype showed some tissue regression at the start of the experiment. However, these control clones fully recovered after one day.

The initial cellular response of marine sponges to thermal stress is often geared towards preserving cellular functions, allowing the holobiont to persist within the environment. Frequently, this involves the upregulation of the proteolytic system to degrade damaged proteins, as found in the sponges *Spongia officinalis* and *Rhopaloeides odorabile* [49, 52]. In *I. basta*, the first temperature induced changes in the transcriptomes occurred between 0h and 2 days, with 17 (thin) and 43 (thick) significantly upregulated genes identified at day 2 in thermally stressed samples compared to control samples (**Figure S2A**). At this time point, the thin morphotype showed an upregulation of stress-response related genes, including heat shock protein 90 (Hsp90) and Wnt-5a. These genes play an important role in directing degraded proteins to proteolysis (Hsp90 [53]) and in tissue regeneration, development and wound healing (Wnt [54]). Similarly, the thick morphotype showed an upregulation of Hsp90 co-chaperones, as well as the ubiquitin-conjugating enzyme, indicating the full activation of the proteolytic system to degrade damaged proteins across morphotypes [55], aiding *I. basta* to maintain cellular functions at the onset of thermal stress.

While both morphotypes showed a similar initial molecular response to heat stress, the subsequent molecular response (from day 4 onwards) reflected the observed difference in their morphological response. For example, after 8 days, the thick morphotype differentially expressed a total of 5,444 genes, compared to 3,784 genes in the thin morphotype (**Figure S2, Table S4**). Among genes that were differentially expressed in the thick morphotype were those linked to metabolic failure, including the downregulation of Hsp70 [56]. However, it remains challenging to elucidate what drives the faster heat stress response of the thick morphotype, with >5,000 genes differentially expressed throughout the experiment. Furthermore, physiological differences between the morphotypes and varying environmental conditions experienced *in situ* may influence the sponge’s stress response [57]. Thus, we suggest future work to focus on characterising these variables to elucidate if and how they impact response to ocean warming.

### Identification of commonly up- and downregulated gene ontology terms in response to heat stress

Due to the observed individual effects and the differences in the molecular response between morphotypes, we focused on how heat stress impacted the biological processes of *I. basta* in general (i.e. in all individuals). To remove the individual effect from our analysis and focus on the effect of temperature alone (**Figure S1**), weighted gene correlation network analyses (WGCNA) were performed. WGCNA separates groups of genes that change their expression in response to temperature from genes that change their expression in response to individual and/or time point (see **Note S1**). A total of 4 (thin) and 7 (thick) modules (i.e. groups of genes) were found to be solely correlated to temperature, each containing between 284 and 5,190 (thin) and 324 and 4,779 (thick) genes. A gene ontology (GO) enrichment analysis was run on these 11 modules to identify GO terms that were significantly up- or downregulated in both morphotypes. This resulted in a total of 60 commonly upregulated (**Figure 2A**) and 74 downregulated GO terms (**Figure 2B**).

### Ianthella basta undergoes apoptosis after 4 days of heat stress

To prevent or minimize cellular damage at the onset of thermal stress, marine sponges are not only thought to upregulate their proteolytic system, but also their immune response and defence [58]. Similarly, in *I. basta*, numerous GO terms associated with these functions were enriched in heat stressed samples, such as “innate immune response”, “T cell activation”, “positive regulation of I-kappaB kinase/NF-kappaB signaling”, “protein ubiquitination”, and “protein neddylation” (**Figure 2A**, see **Table S5** for the individual genes per GO term). While these functions seem to aid *I. basta* in coping with increasing SST, they ultimately failed to prevent cellular damage and *I. basta* underwent tissue regression after 4 to 8 days (**Figure 1**). Congruently, numerous upregulated GO terms in *I. basta* were pathways related to apoptosis (i.e. programmed cell death; **Figure 2A, Table S5**). Apoptosis is tightly linked to oxidative stress [59] and has been proposed as one of the cellular mechanisms that could drive the breakdown of symbiosis in corals [60, 61]. Similarly, apoptosis may drive dysbiosis in marine sponges and characterising the mechanism and onset of apoptosis may aid in understanding the breakdown of the sponge holobiont. Generally, apoptosis can occur through two pathways: the intrinsic or mitochondrial pathway and extrinsic or death receptor pathway [62]. Here, we suggest that *I. basta* is undergoing apoptosis through the intrinsic pathway. Firstly, the GO term “positive regulation of calcium ion transport” was upregulated. Ca^2+^ is an important signal in the pro-apoptotic transition of mitochondria [63, 64]. An overload of Ca^2+^ can induce the swelling of mitochondria, resulting in the rupture of the outer membrane and the release of apoptotic factors [65]. Damaged mitochondria can be controlled through autophagic degradation, of which related GO terms were upregulated in *I. basta* (i.e. “autophagy of mitochondria” and “positive regulation of autophagy of mitochondrion in response to mitochondrial depolarization”). If, however, mitochondria rupture and release apoptotic factors, a signalling cascade will be initiated that leads to apoptosis. This signalling pathway involves the proteins Caspases 3 and 7 [64], which were significantly upregulated in *I. basta* after 4 days of heat stress.

TEM images from control and heat stressed samples from one thin individual from T = 8 days provided further support for apoptosis (see Supplementary Methods and **Figure S4**). These images show putative late stages of apoptosis, indicating that apoptosis occurred between 2 to 8 days of heat stress.

### Heat stress further reduces Ianthella basta’s ability to mitigate tissue damage and pump water

To mitigate cellular damage, sponges can rapidly regenerate their tissue [66-68]. Sponges transform epithelial cells into mesenchymal stem cells which can subsequently differentiate into a variety of cell types [69]. In *I. basta*, heat stress induced the downregulation of the GO term “positive regulation of epithelial to mesenchymal transition” after two to four days (**Figure 2B, Table S6**), suggesting a reduced ability of the sponge to regenerate its tissue after undergoing apoptosis. Furthermore, under heat stress, *I. basta*’s ability to maintain overall tissue integrity decreased, indicated by a downregulation of the GO terms “actin crosslink formation” and “heterophilic cell-cell adhesion via plasma membrane cell adhesion molecules”, amongst others (**Figure 2B, Table S6**). Notably, genes involved in the maintenance of tissue integrity also had some of the highest log2 fold change in the differential gene abundance analysis (e.g. actin, actin binding proteins, and collagen alpha chain proteins). Heat stress also appeared to hamper the pumping capacity of *I. basta* (i.e. downregulation of “regulation of cilium beat frequency”, “epithelial cilium movement involved in extracellular fluid movement”, and “motile cilium assembly”). If *I. basta* stops or reduces its pumping rate, nutrient uptake could be impaired, further destabilising the holobiont. Moreover, the sponge tissue can become anoxic, impacting metabolic interactions at the microscale [70], potentially disproportionately affecting the thick morphotype [57].

### Thermal stress reduces the ability of the host to recognise symbionts

To further identify molecular mechanisms contributing to dysbiosis in marine sponges, we characterised the effect of thermal stress on *I. basta*’s immune system. Sponges have a well-developed innate immune system, comprising a wide range of pattern-recognition receptors (PRRs) designed to recognise microbes and pathogens via microbial-associated molecular patterns (MAMPs) [50, 71]. In *I. basta*, two PRRs, the scavenger receptor cysteine-rich (SRCR) family members and G protein-coupled receptors (GPCRs, belonging to the adhesion family), were downregulation in heat stressed samples after 4 days (**Table S4**), suggesting a reduced capacity of the sponge to distinguish between beneficial microbes and pathogens. In conjunction, mechanisms that aid in the defence against extracellular bacterial infections were upregulated (GO term: “interleukin-17-mediated signaling pathway”, **Figure 2, Table S5**).

### Thermal stress causes significant changes in the microbial community after 8 days

To reveal if *I. basta*’s stress response and reduced ability to differentiate between microbes correlated to changes in the microbial community composition, we completed 16S rRNA gene amplicon sequencing on all samples. A total of 2,161,768 reads passed quality filtering, in which 756 amplicon sequence variants (ASVs) were identified (**Table S6**). These ASVs were classified into 24 bacterial phyla and 2 archaeal phyla. Consistent with previous work on *I. basta*’s microbiome [27, 72], the microbial community of healthy individuals was dominated by three microbial ASVs: a Nitrososphaeria (ASV_2919 thin, ASV_2919 thick), Gammaproteobacteria (ASV_1194 thin, ASV_496 thick), and Alphaproteobacteria (ASV_572 thin, ASV_1253 thick; **Figure 3**). Each of these ASVs displayed >99.20% sequence identity when mapped against the respective genomes of the three dominant symbionts, ‘*Ca*. Nitrosospongia ianthellae’, ‘*Ca*. Taurinisymbium ianthellae’ and ‘*Ca*. Luteria ianthellae’, respectively, which have previously been recovered and functionally annotated [25-27, 73].

**Figure 3.**
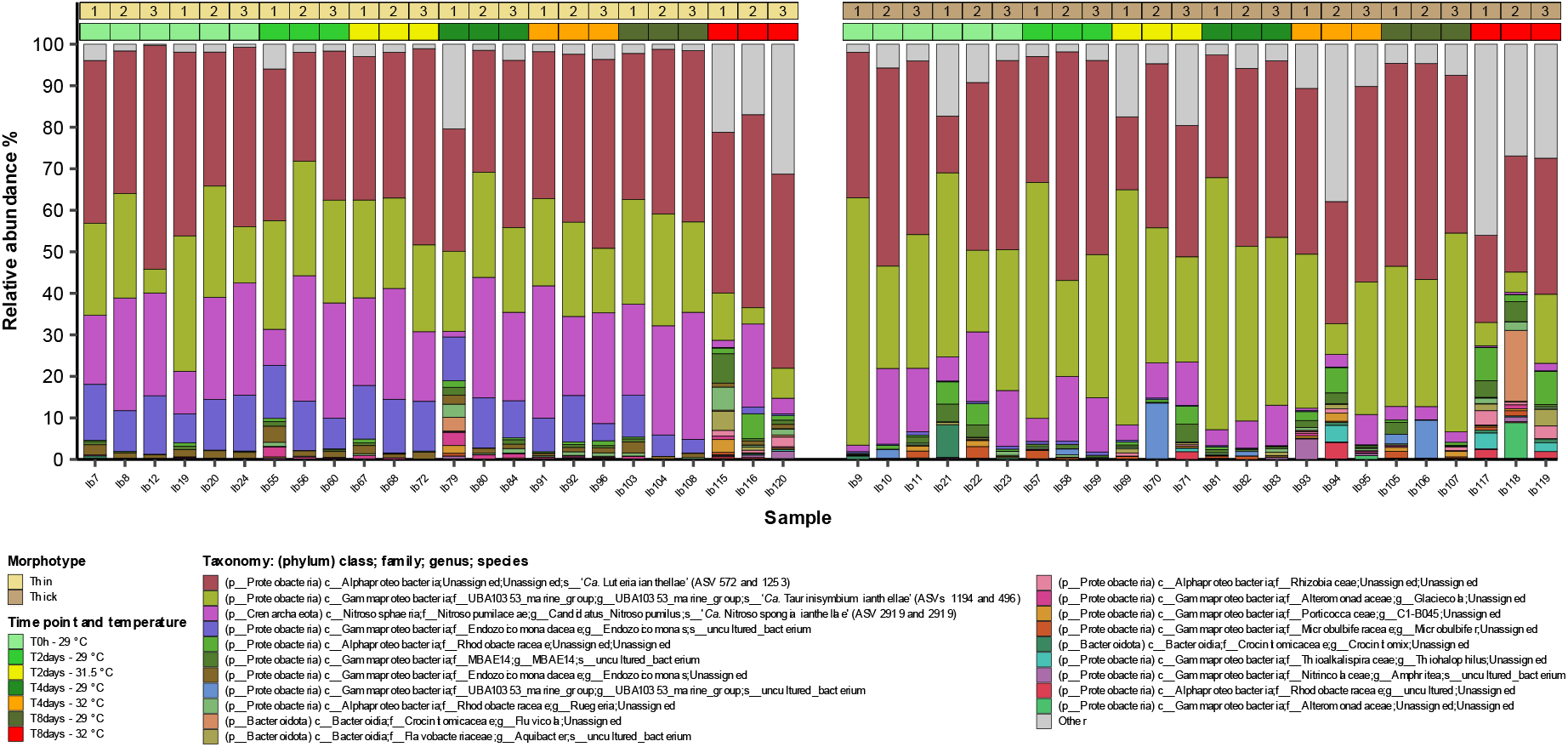
Barplot displaying the 20 most abundant microbial species in *I. basta* individuals with a thin (left) and thick (right) morphotype in control and treatment sample at T = 0h, T= 2 days, T = 4 days, and T = 8 days. The lowest resolved taxonomy is shown, and for simplicity is displayed as phylum, class, family, genus, and species. Ib: *Ianthella basta*, followed by sample number. The number displayed on top of the morphotype bar represents the individual number.

Shifts in the microbial community of sponges have often been observed in response to heat stress and frequently occur when the sponge starts to show substantial stress (e.g. necrosis) [21, 23, 30, 74], although exceptions exist [22, 24]. In *I. basta*, the shift in the microbial community seemed to follow the physiological stress response of the host. In the thin morphotype of *I. basta*, for example, the microbiome was highly stable up to day 4 of the experiment and only became significantly different after 8 days (PERMANOVA, p < 0.05, **Figures 3 and S6**). This difference was predominantly driven by a decrease in the relative abundance of the Nitrososphaeria (SIMPER analysis, contribution of 2.49%), which may be caused by the reduced ability of the host to recognise the symbiont, by the observed apoptosis and breakdown of the host tissue, or by the change in the local nutrient and oxygen conditions associated with a reduced pumping rate. In contrast, the thick morphotype already showed (a non-significant) decrease in the relative abundance of the three dominant symbionts after 4 days, matching its faster morphological response (**Figure 3, Figure S5**). After 8 days, the microbial community became significantly different between control and treatment samples (PERMANOVA, p < 0.05, **Figures 3 and S5**), driven by a decrease in the relative abundance of the dominant Gammaproteobacterium (SIMPER analysis, contribution of 3.07%). The relative abundance of the dominant Alphaproteobacterium did not change significantly for either of the morphotypes. Changes in the relative abundance of the dominant symbionts was further supported by visual observations of FISH images (see Supplementary Methods and **Figure S6**).

The reduction in the abundance of the dominant symbionts at day 8 co-occurred with the introduction of 414 unique microbial ASVs (**Figure 3, Table S7**), of which some displayed distinct morphologies (**Figure S6**). Most of these introduced ASVs belonged to the taxa Bacteroidota (22% of the ASVs, e.g. orders Chitinophagales and Flavobacteriales), Planctomycetota (21%, e.g. orders OM190 and Phycisphaerales), and Alpha- and Gammaproteobacteria (20% and 16%, respectively, e.g. orders Cellvibrionales and Caulobacterales), thus showing similar taxonomic shifts to previous reports on the microbial community of necrotic sponges [21, 23, 28]. Members of the introduced taxa, such as the Bacteroidota (e.g. Flavobacteriaceae) and Gammaproteobacteria (e.g. Alteromonadaceae), are established opportunists in the degradation of organic matter [75, 76] and are likely attracted by the release of nutrients from degrading sponge tissue. Their introduction may be facilitated by the overall breakdown of the sponge host, including its decreased ability to distinguish between microbes and pathogens, as well as the reduced abundance of the dominant symbionts, creating available niche space.

### Thermal stress results in the loss of symbiotic functions

The health of *I. basta* is intimately linked to its symbiosis with microorganisms, which are involved in the removal of waste products (i.e. ammonia), fixation of carbon, and the provision of vitamins and essential amino acids [25, 27, 73]. When, in response to heat stress, key symbionts reduce in abundance, symbiotic functions may be lost, especially in low microbial diversity sponges where functional redundancy is low and other microbes cannot easily replace the function [77]. To investigate whether the observed changes in the microbial community of *I. basta* translated into a loss of symbiotic functions, we processed all samples from day 4 and day 8 for metaproteomics. A total of 4,490 and 4,431 actively expressed proteins were retrieved for the microbial enriched protein fraction of the thin and thick morphotype, respectively. Of these proteins, 2,600 and 2,571 were functionally annotated, either by the (KEGG) Orthologies (KOs) or PFAM database (**Table S8**). A total of 72 (thin) and 80 (thick) were captured in the genomes of *I. basta*’s symbionts, which have been previously recovered and annotated [27].

The dominant archaea in *I. basta* plays a key role in the recycling of nitrogenous waste products (i.e. ammonium, urea) and autotrophic carbon fixation, potentially benefitting the sponge host [25, 27, 73]. Although we did not recover ammonia monooxygenase (amoA – the protein responsible for the oxidation of toxic ammonia) in the metaproteomic data, likely because it is membrane bound [78], the reduced abundance of the ammonia-oxidizing-archaeaon could result in an accumulation of nitrogenous waste products. This accumulation would be further exacerbated by any reduction in the pumping rates of the sponge host. Other key processes of removing and transforming nitrogen compounds from the sponge host, performed by the dominant archaea and gammaproteobacterium [27], were also lost in response to heat stress. For instance, *nirK*, the enzyme needed to perform the first step of the reduction of nitrite to dinitrogen through nitric oxide (i.e. denitrification) was absent in day 8 heat stressed samples of both morphotypes, while present in control samples (thin: sequence_2220078, thick: sequence_476815, **Table S8**). To supplement the metaproteomic data, we visualised the effect of heat stress on the ability of the dominant archaea to fix dissolved inorganic carbon (DIC), by combining advanced stable isotope tracing with fluorescence and chemical imaging (see Supplementary Methods). ^13^C-DIC incubations suggested a reduction in the assimilation of inorganic carbon in *I. basta* after 8 days of heat stress, coinciding with the reduced abundance of the archaea (**Figure S7**). However, due to the low number of archaea in the heat stress sample present and measured, we were unable to draw statistically supported conclusions on differences in single cell activities over incubation time. Given the already reduced archaeal abundance at day 8, heat stress may have affected symbiont activity earlier. The sponge microbiome has also been suggested to contribute to the production of essential branched-chain amino-acids (BCAAs, e.g. leucine, isoleucine, and valine) and B-vitamins [79, 80], something the host cannot produce itself, which has also been observed in *I. basta* [73]. Heat stress can disrupt this nutritional interdependence by causing a reduction in BCAA biosynthesis and transport [21, 30, 81]. After 4 days of heat stress, we observed a reduced expression of genes involved in the biosynthesis of BCAAs (i.e. LeuA, ilvA, and ilvB, **Table S8**). Furthermore, after 8 days of heat stress, the Gammaproteobacterial symbiont of the thin *I. basta* morphotype showed no expression of the livK gene (sequence_1700135, **Table S8**), the substrate binding unit of the LIV-I BCAA system. LivK was also absent in day 8 heat stressed samples from the thick morphotype (sequence_3584319). Notably, in the host transcriptome of the thick *I. basta* morphotype, genes assigned to the GO terms “basic amino acid transport” and “cobalamin transport” were downregulated after 4 days of heat stress (**Table S4**), suggesting not only a reduced uptake of amino-acids, but also of essential B-vitamins.

Analysis of the metaproteomic data further provided insight into putative mechanisms that may cause dysbiosis. To persist within the sponge tissue, symbionts can attach directly to the host or to other microbes that are attached to the host tissue. In *I. basta*, several proteins have previously been identified that mediate this cell-cell adhesion and biofilm formation, including cadherin domains and fibronectin binding proteins [27]. Interestingly, heat stress caused a loss of these attachment proteins, with cadherin(-like) domains, as well as fibronectin III domains being absent in heat stressed samples at day 8 in the thin morphotype (sequences 151070, 2336744, 3714211, and 1867271, **Table S8**). Thus, while it is tempting to speculate that the reduced abundance of the dominant symbionts may be caused by the reduced ability of the sponge host to distinguish between symbionts and non-symbiotic microbes, or by the observed apoptosis and breakdown of the host tissue, it could also be mediated by a reduced expression of attachment proteins.

## Conclusion and outlook

Climate change driven ocean warming has caused significant declines in vital reef organisms, such as corals and sponges. To fully understand the effects of increasing SST, we present the integrated response of the model sponge *I. basta* to heat stress and provide key insights into the response of both the host and symbionts (**Figure 4**). *I. basta* appears to maintain cellular homeostasis at the onset of thermal stress, through the upregulation of immune related genes and genes that aid in the degradation of damaged proteins. However, the upregulation of apoptotic pathways, downregulation of tissue integrity pathways and pumping capacity, and the observed tissue regression after 4 to 8 days of heat stress, indicates that *I. basta* ultimately fails to cope with seawater temperatures (+ 1.5 C) predicted to occur within this century [82]. Moreover, after 4 days, the sponge showed a reduced expression of PRRs to distinguish between symbionts and opportunistic microbes. Congruently, a reduced relative abundance of the three dominant symbionts occurred from day 4 onwards, which was followed by an introduction of opportunistic microbes. The observed reduction in the relative abundance of the dominant symbionts resulted in a relative reduction of detected symbiotic functions and attachment proteins, further disrupting the interdependence between the sponge host and its symbionts. As such, this work provides new insights into the heat stress response of marine sponges and highlights how future SST can impact host-microbe interaction in these vital organisms.

**Figure 4.**
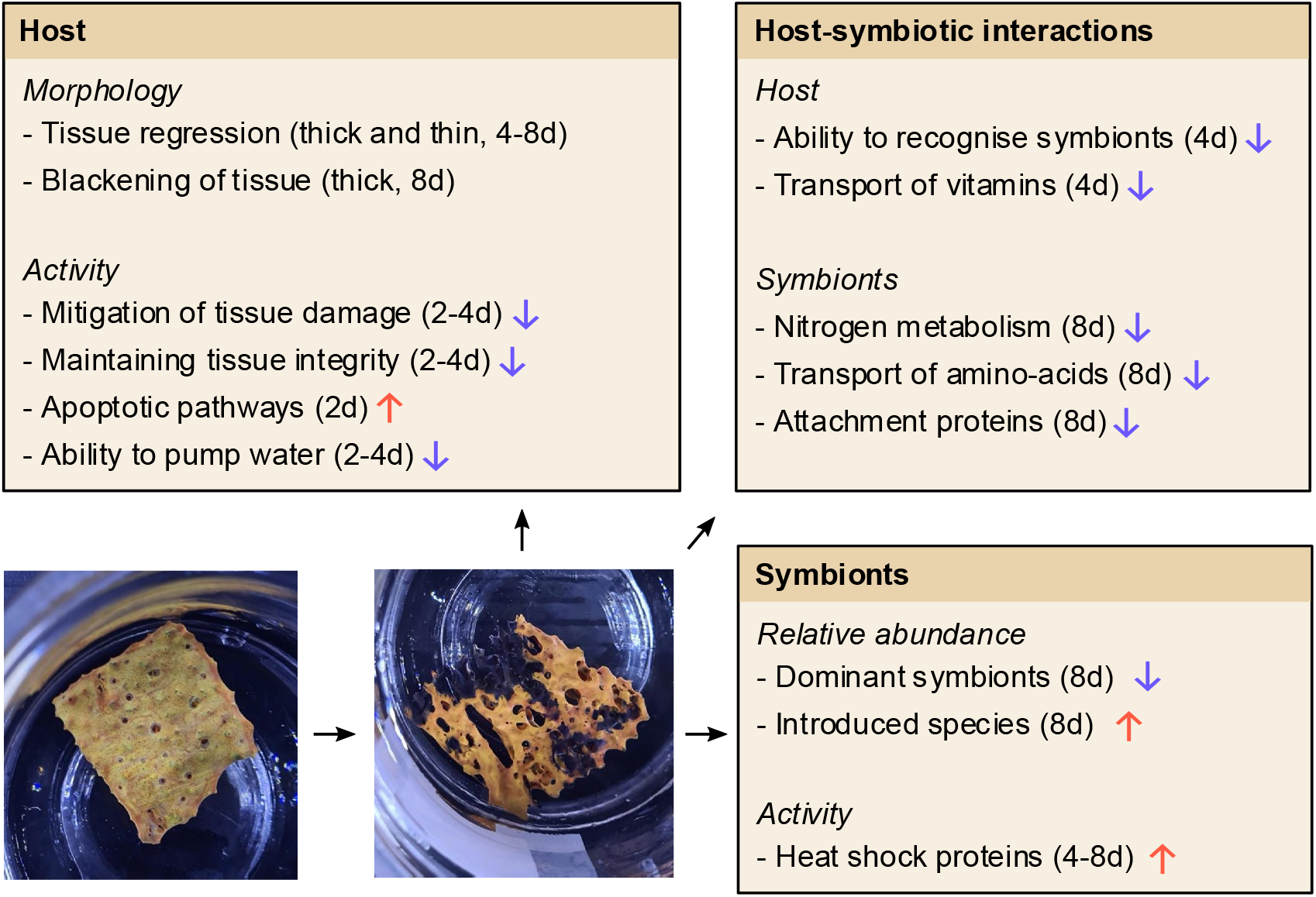
Overview figure of the holobiont response of *Ianthella basta* to heat stress, with arrows pointing up indicating upregulation of the respective function and arrow pointing down indicating downregulation. The time of onset of the stress response is also shown.

## Acknowledgements

We would like to thank Stefan Schuster, Sarah Bosiljcic and Stefanie Imminger from the Centre of Microbiology and Environmental Systems Science for their contribution with sample processing. We also acknowledge the technical support of the Histology Facility at Vienna BioCenter Core Facilities (VBCF), member of the Vienna BioCenter (VBC), Austria, and especially thank Mihaela Grivej for her help. We also thank Emmanuelle Botté for her help with sample collection.

## Contributions

NSW, JPE, and LR conceptualised and designed the study; JPE, LR, and HL ran the experiment and all authors contributed to data generation or interpretation. JPE wrote the manuscript with contributions from all authors.

## Data availability

The mass spectrometry proteomics data have been deposited to the ProteomeXchange Consortium via the PRIDE partner repository. RNA and 16S rRNA gene amplicon sequencing data have been deposited at the Sequence Read Archive.

## References

1. De’Ath, G., et al., The 27–year decline of coral cover on the Great Barrier Reef and its causes. Proc Natl Acad Sci, 2012. 109(44): p. 17995–17999.

2. Webster, N.S., Sponge disease: a global threat? Env Microbiol, 2007. 9(6): p. 1363–1375.

3. Hughes, T.P., et al., Global warming and recurrent mass bleaching of corals. Nature, 2017. 543(7645): p. 373–377.

4. Perkins, N.R., et al., Bleaching in sponges on temperate mesophotic reefs observed following marine heatwave events. Clim Chang Ecol, 2022. 3: p. 100046.

5. IPCC, Summary for policymakers, in Climate Change 2021: The Physical Science Basis. Contribution of Working Group I to the Sixth Assessment Report of the Intergovernmental Panel on Climate Change, P.Z.V. Masson-Delmotte, A. Pirani, S.L. Connors, C. Péan, S. Berger, N. Caud, Y. Chen, L. Goldfarb, M.I. Gomis, M. and K.L. Huang, E. Lonnoy, J.B.R. Matthews, T.K. Maycock, T. Waterfield, O. Yelekçi, R. Yu, and B. Zhou, Editors. 2021, Cambridge University Press.

6. Hoegh-Guldberg, O., et al., Coral reefs under rapid climate change and ocean acidification. Science, 2007. 318(5857): p. 1737–1742.

7. Cziesielski, M.J., S. Schmidt-Roach, and M. Aranda, The past, present, and future of coral heat stress studies. Ecol Evol, 2019. 9(17): p. 10055–10066.

8. Webster, N.S., et al., Host-associated coral reef microbes respond to the cumulative pressures of ocean warming and ocean acidification. Sci Rep, 2016. 6(1): p. 19324.

9. Savary, R., et al., Fast and pervasive transcriptomic resilience and acclimation of extremely heat-tolerant coral holobionts from the northern Red Sea. Proc Natl Acad Sci, 2021. 118(19): p. e2023298118.

10. Quigley, K.M. and M.J.H. van Oppen, Predictive models for the selection of thermally tolerant corals based on offspring survival. Nat Commun, 2022. 13(1): p. 1543.

11. Przeslawski, R., et al., Beyond corals and fish: the effects of climate change on noncoral benthic invertebrates of tropical reefs. Glob Change Biol, 2008. 14(12): p. 2773–2795.

12. Wolfe, K., et al., Priority species to support the functional integrity of coral reefs, in Oceanography and Marine Biology. 2020, Taylor & Francis.

13. Bell, J.J., The functional roles of marine sponges. Estuar Coast Shelf Sci, 2008. 79(3): p. 341–353.

14. Bell, J.J., et al., Interocean patterns in shallow water sponge assemblage structure and function. Biol Rev, 2020. 95(6): p. 1720–1758.

15. Bell, J.J., et al., Sponges to be winners under near-future climate scenarios. Biosci, 2018. 68(12): p. 955–968.

16. de Goeij, J.M., et al., Surviving in a marine desert: the sponge loop retains resources within coral reefs. Science, 2013. 342(6154): p. 108–110.

17. Pita, L., et al., The sponge holobiont in a changing ocean: from microbes to ecosystems. Microbiome, 2018. 6(1): p. 46.

18. Robbins, S., et al., A genomic view of the microbiome of coral reef demosponges. ISME J, 2021: p. 1–14.

19. Hudspith, M., et al., Subcellular view of host–microbiome nutrient exchange in sponges: insights into the ecological success of an early metazoan–microbe symbiosis. Microbiome, 2021. 9(1): p. 1–15.

20. Wilkinson, C.R., Net primary productivity in coral reef sponges. Science, 1983. 219(4583): p. 410–412.

21. Fan, L., et al., Marine microbial symbiosis heats up: the phylogenetic and functional response of a sponge holobiont to thermal stress. ISME J, 2013. 7(5): p. 991–1002.

22. Ramsby, B.D., et al., Elevated seawater temperature disrupts the microbiome of an ecologically important bioeroding sponge. Mol Ecol, 2018. 27(8): p. 2124–2137.

23. Simister, R., et al., Thermal stress responses in the bacterial biosphere of the Great Barrier Reef sponge, Rhopaloeides odorabile. Environ Microbiol, 2012. 14(12): p. 3232–3246.

24. Webster, N.S., R.E. Cobb, and A.P. Negri, Temperature thresholds for bacterial symbiosis with a sponge. ISME J, 2008. 2(8): p. 830.

25. Moeller, F.U., et al., Characterization of a thaumarchaeal symbiont that drives incomplete nitrification in the tropical sponge Ianthella basta. Environ Microbiol, 2019: p. 527234.

26. Moeller, F.U., et al., Taurine as a key intermediate for host-symbiont interaction in the tropical sponge Ianthella basta. ISME J, 2023: p. 1–16.

27. Engelberts, J.P., et al., Metabolic reconstruction of the near complete microbiome of the model sponge Ianthella basta. Env Microbiol, 2022. 25(3): p. 646–660.

28. Webster, N.S., R.E. Cobb, and A.P. Negri, Temperature thresholds for bacterial symbiosis with a sponge. ISME J, 2008. 2(8): p. 830–842.

29. Luter, H.M., S. Whalan, and N.S. Webster, Thermal and sedimentation stress are unlikely causes of brown spot syndrome in the coral reef sponge, Ianthella basta. PLoS One, 2012. 7(6).

30. Botté, E.S., et al., Future ocean conditions induce necrosis, microbial dysbiosis and nutrient cycling imbalance in the reef sponge Stylissa flabelliformis. ISME Commun, 2023. 3(1).

31. Meyer, E., G. Aglyamova, and M. Matz, Profiling gene expression responses of coral larvae (Acropora millepora) to elevated temperature and settlement inducers using a novel RNA-Seq procedure. Mol Ecol, 2011. 20(17): p. 3599–3616.

32. Lohman, B.K., J.N. Weber, and D.I. Bolnick, Evaluation of TagSeq, a reliable low-cost alternative for RNA seq. Mol Ecol Resour, 2016. 16(6): p. 1315–1321.

33. Martin, M., Cutadapt removes adapter sequences from high-throughput sequencing reads. EMBnet j, 2011. 17(1): p. 10–12.

34. Langmead, B. and S.L. Salzberg, Fast gapped-read alignment with Bowtie 2. Nat Methods, 2012. 9(4): p. 357–359.

35. Love, M.I., W. Huber, and S. Anders, Moderated estimation of fold change and dispersion for RNA-seq data with DESeq2. Genome Biol, 2014. 15(12): p. 1–21.

36. R Core Team, R: A language and environment for statistical computing. 2013.

37. Langfelder, P. and S. Horvath, WGCNA: an R package for weighted correlation network analysis. BMC bioinform, 2008. 9(1): p. 1–13.

38. Alexa, A. and J. Rahnenfuhrer, topGO: enrichment analysis for gene ontology. R package version, 2010. 2(0): p. 2010.

39. Parada, A.E., D.M. Needham, and J.A. Fuhrman, Every base matters: assessing small subunit rRNA primers for marine microbiomes with mock communities, time series and global field samples. Env Microbiol, 2016. 18(5): p. 1403–1414.

40. Apprill, A., et al., Minor revision to V4 region SSU rRNA 806R gene primer greatly increases detection of SAR11 bacterioplankton. Aquat Microb Ecol, 2015. 75(2): p. 129–137.

41. Bolger, A.M., M. Lohse, and B. Usadel, Trimmomatic: a flexible trimmer for Illumina sequence data. Bioinform, 2014. 30(15): p. 2114–2120.

42. Bolyen, E., et al., Reproducible, interactive, scalable and extensible microbiome data science using QIIME 2. Nat Biotechnol, 2019. 37(8): p. 852–857.

43. Callahan, B.J., et al., DADA2: high-resolution sample inference from Illumina amplicon data. Nat Methods, 2016. 13(7): p. 581.

44. Clarke, K. and R. Gorley, Getting started with PRIMER v7. PRIMER-E: Plymouth, Plymouth Marine Laboratory, 2015. 20.

45. Kenny, N.J., et al., Towards the identification of ancestrally shared regenerative mechanisms across the Metazoa: A transcriptomic case study in the Demosponge Halisarca caerulea. Mar Genomics, 2018. 37: p. 135–147.

46. Luter, H.M., et al., Gene correlation networks reveal the transcriptomic response to elevated nitrogen in a photosynthetic sponge. Mol Ecol, 2020. 29(8): p. 1452–1462.

47. Strehlow, B.W., et al., Novel reference transcriptomes for the sponges Carteriospongia foliascens and Cliona orientalis and associated algal symbiont Gerakladium endoclionum. Coral Reefs, 2021. 40(1): p. 9–13.

48. Desplat, Y., J.F. Warner, and J.V. Lopez, Holo-transcriptome sequences from the tropical marine sponge Cinachyrella alloclada. J Hered, 2021.

49. Koutsouveli, V., et al., Gearing up for warmer times: transcriptomic response of Spongia officinalis to elevated temperatures reveals recruited mechanisms and potential for resilience. Front Mar Sci, 2020. 6.

50. Pita, L., et al., Differential expression of immune receptors in two marine sponges upon exposure to microbial-associated molecular patterns. Sci Rep, 2018. 8(1): p. 16081.

51. Posadas, N., et al., Microbiome diversity and host immune functions influence survivorship of sponge holobionts under future ocean conditions. ISME J, 2022. 16(1): p. 58–67.

52. Pantile, R. and N. Webster, Strict thermal threshold identified by quantitative PCR in the sponge Rhopaloeides odorabile. Mar Ecol Prog Ser, 2011. 431: p. 97–105.

53. Richter, K., M. Haslbeck, and J. Buchner, The heat shock response: Life on the verge of death. Mol Cell, 2010. 40(2): p. 253–266.

54. Wu, Y.-C., et al., Wounding response in Porifera (sponges) activates ancestral signaling cascades involved in animal healing, regeneration, and cancer. Sci Rep, 2022. 12(1): p. 1307.

55. Fernández-Fernández, M.R., et al., Hsp70–a master regulator in protein degradation. FEBS letters, 2017. 591(17): p. 2648–2660.

56. López-legentil, S., et al., Bleaching and stress in coral reef ecosystems: hsp70 expression by the giant barrel sponge Xestospongia muta. Mol Ecol, 2008. 17(7): p. 1840–1849.

57. Massaro, A.J., et al., Behavioral and morphological changes caused by thermal stress in the Great Barrier Reef sponge Rhopaloeides odorabile. J Exp Mar Biol Ecol, 2012. 416-417: p. 55–60.

58. Guzman, C. and C. Conaco, Gene expression dynamics accompanying the sponge thermal stress response. PloS one, 2016. 11(10): p. e0165368.

59. Lesser, M.P., Oxidative stress in marine environments: Biochemistry and Physiological Ecology. Annu Rev Physiol, 2006. 68(1): p. 253–278.

60. Ainsworth, T.D., et al., Early cellular changes are indicators of pre-bleaching thermal stress in the coral host. J Exp Mar Biol Ecol, 2008. 364(2): p. 63–71.

61. Lesser, M.P., Oxidative stress causes coral bleaching during exposure to elevated temperatures. Coral reefs, 1997. 16(3): p. 187–192.

62. Elmore, S., Apoptosis: A review of programmed cell death. Toxicol Pathol, 2007. 35(4): p. 495–516.

63. Kroemer, G. and J.C. Reed, Mitochondrial control of cell death. Nat Med, 2000. 6(5): p. 513–519.

64. Giorgi, C., et al., Mitochondrial Ca2+ and apoptosis. Cell Calcium, 2012. 52(1): p. 36–43.

65. Rizzuto, R., et al., Ca2+ signaling, mitochondria and cell death. Cur Mol Med, 2008. 8(2): p. 119–130.

66. Luter, H.M., S. Whalan, and N.S. Webster, The marine sponge Ianthella basta can recover from stress-induced tissue regression. Hydrobiologia, 2012. 687(1): p. 227–235.

67. Ayling, A.L., Growth and regeneration rates in thinly encrusting demospongiae from temperate waters. Biol Bull, 1983. 165(2): p. 343–352.

68. Wulff, J., Regeneration of sponges in ecological context: is regeneration an integral part of life history and morphological strategies? Integr Comp Biol, 2010. 50(4): p. 494–505.

69. Ereskovsky, A.V., et al., Transdifferentiation and mesenchymal-to-epithelial transition during regeneration in Demospongiae (Porifera). J Exp Zool, 2020. 334(1): p. 37–58.

70. Hoffmann, F., et al., An anaerobic world in sponges. Geomicrobiol J, 2005. 22(1-2): p. 1–10.

71. Hentschel, U., et al., Genomic insights into the marine sponge microbiome. Nat Rev Microbiol, 2012. 10(9): p. 641–654.

72. Luter, H.M., S. Whalan, and N.S. Webster, Exploring the role of microorganisms in the disease-like syndrome affecting the sponge Ianthella basta. Appl Environ Microbiol, 2010. 76(17): p. 5736–5744.

73. Glasl, B., et al., Branched-chain amino acid assimilation enables mixotrophy of ammonia-oxidizing archaeal sponge symbionts. Sci Adv, 2026. 12(36): p. eaef9450.

74. Núñez-Pons, L., et al., Too hot for my bugs: mediterranean heatwave disrupts associated microbiomes in the sponge Petrosia ficiformis. Environmental Microbiome, 2025.

75. Edwards, J.L., et al., Identification of carbohydrate metabolism genes in the metagenome of a marine biofilm community shown to be dominated by Gammaproteobacteria and Bacteroidetes. Genes, 2010. 1(3): p. 371–384.

76. Francis, B., et al., North Sea spring bloom-associated Gammaproteobacteria fill diverse heterotrophic niches. Environ Microbiome, 2021. 16(1): p. 15.

77. Cárdenas, A., et al., Greater functional diversity and redundancy of coral endolithic microbiomes align with lower coral bleaching susceptibility. ISME J, 2022.

78. Hyman, M.R. and D. Arp, 14C2H2-and 14CO2-labeling studies of the de novo synthesis of polypeptides by Nitrosomonas europaea during recovery from acetylene and light inactivation of ammonia monooxygenase. Journal of Biological Chemistry, 1992. 267(3): p. 1534–1545.

79. Song, H., O.H. Hewitt, and S.M. Degnan, Arginine biosynthesis by a bacterial symbiont enables nitric oxide production and facilitates larval settlement in the marine-sponge host. Curr Biol, 2021. 31(2): p. 433-437. e3.

80. Engelberts, J.P., et al., Characterization of a sponge microbiome using an integrative genome-centric approach. ISME J, 2020. 15(5): p. 1100–1110.

81. Chai, G., J. Li, and Z. Li, The interactive effects of ocean acidification and warming on bioeroding sponge Spheciospongia vesparium microbiome indicated by metatranscriptomics. Microbiological Research, 2024. 278: p. 127542.

82. Pörtner, H.-O., et al., Climate change 2022: Impacts, adaptation and vulnerability. 2022.

