## Supplementary Information for "Thermal stress disrupts microbial symbiosis and host activity in the model sponge *Ianthella basta*"

#### Supplementary Methods

*Host reference transcriptome: RNA extraction, sequencing, de novo assembly, and annotation*

To generate a host reference transcriptome for both morphotypes of *I. basta* (thin and thick), total RNA was extracted from nine samples per morphotype, covering all individuals, timepoints, and temperature conditions at least once to capture the full range of gene expression responses. Tissue of each sample was ground on liquid nitrogen after which RNA was extracted with TRIzol reagent (Thermo Fisher Scientific) following the manufacturer's instructions, using a mix of 1.4- and 0.1-mm beads (MP Biomedicals) for the homogenization of samples and an additional bead-beating step at 2500 rpm for 40 s. Extracted RNA was purified using the PureLink RNA Mini Kit (Ambion) and further treated with PureLink DNase to remove genomic DNA. The quantity of the total RNA was measured using Qubit. Equal amounts of RNA were combined for samples from the same morphotype to a final concentration of 100 ng/µl. The resulting two samples were sent to the Australian Centre for Ecogenomics, Brisbane, for sequencing. Here, libraries were prepared with 400 ng of total RNA using the lllumina Stranded Total RNA Prep with Ribo-Zero Plus (Illumina #20040529), following the manufacturer's protocol. Samples were indexed with the IDT for Illumina RNA UD Indexes. Each library was quantified, and quality control was performed using the Quant-iT™ dsDNA HS Assay Kit (Invitrogen) and Agilent D1000 HS tapes (#5067-5582) on the TapeStation 4200 (Agilent #G2991AA), following the manufacturer's instructions. Libraries were pooled in equimolar ratio and sequenced on a NovaSeq6000 (Illumina) using NovaSeq6000 SP kit v1.5, 2 x 150 bp paired end, generating a minimum of 100 million reads per sample.

Adaptor sequences were removed using bcl2fastq v2.20.0.422 and reads were quality filtered using fastx_clipper (http://hannonlab.cshl.edu/fastx_toolkit), retaining reads longer than 50 nucleotides with a minimum quality score of 20 over at least 80% of the read. Reads were subsequently assembled into two reference transcriptomes, one for each morphotype, using Trinity v2.12.0 [1]. Transcriptome completeness was estimated by identifying single-copy orthologs in the transcriptomes, using BUSCO v5.0.0 [2] with the metazoan reference database for conserved genes. In addition to the two separate transcriptomes, a combined reference transcriptome was assembled using all reads. However, the resulting transcriptome was less contiguous and had fewer complete and single-copy orthologues than the individual transcriptomes. Therefore, downstream analyses were performed with the two separate transcriptomes (**Table S2**). Microbial contamination (16S and 23S rRNA gene sequences) of the transcriptomes was assessed with SortMeRNA v4.3.6 [3]. Although microbial contamination was present (thin transcriptome: 2.8% of the reads, thick transcriptome: 17.7% of the reads), all described metabolic processes had a eukaryotic origin (see also **Table S3**).

To functionally annotate the transcriptomes, a set of protein predictions was generated using TransDecoder v5.5.0 (https://github.com/TransDecoder/TransDecoder). Long open reading frames (ORFs) were extracted with the TransDecoder.LongOrfs tool with default parameters. ORFs were then used as queries to perform a blastp v2.9.0+ search against the Uniprot Swissprot database with an E-value cut-off of 1e-5. In addition, conserved Pfam domains were identified within the ORFs based on the Pfam database using HMMER v3.3.1 [4]. Protein predictions were then refined with the TransDecoder.Predict tool, retaining ORFs with blastp and Pfam hits. Furthermore, functional annotations were obtained for each Trinity predicted isoform (~gene) based on homology to proteins in the Swissprot database. To identify homologs in Swissprot, both blastp (for predicted proteins) and blastx (for transcript nucleotide sequences) were used to perform searches, retaining up to 5 hits per query and using an E-value cut-off of 1e-4. Detailed information for all blast hits including gene ontology terms, KEGG identifiers, and gene/protein names was subsequently retrieved using the Uniprot mapping service. For each isoform, a single annotation was chosen as that with the lowest E-value across both blastp and blastx searches. Finally, KEGG identifiers were converted to KO numbers using the KEGG api accessed via the KEGGREST v1.32.0 package in R v4.2.1 [5] (**Table S3**).

*Fluorescence in situ hybridisation (FISH) and transmission electron microscopy (TEM)*

Four samples from T = 8 days (one control and one treatment sample per morphotype) were further processed for FISH to visually inspect changes in microbial abundance and spatial organisation, following the protocol described in Engelberts et al. [49]. In short, samples were sectioned to 5 µm on a Thermofisher Cryostar NX70 cryostat after washing in 1x PBS and embedded in embedding medium (1:1, Surgipath FSC 22 Frozen Section Embedding Medium). Sections were subsequently processed for FISH, visualising all bacteria with the EUB338 I-III probe mix [50] and the dominant ammonia-oxidizing-archaeaon with the ARCH915 probe [25, 51]. The dominant Alpha- and Gammaproteobacterial symbionts of *I. basta* were further visualised using previously validated symbiont specific probes termed AlfD729 and GamD1137, respectively [26].

The same thin individual that was processed for FISH was also processed for TEM imaging to map changes in host tissue morphology at the microscale. Samples were post-fixed in 1% osmium tetroxide in sodium cacodylate, stained with 1% uranyl acetate, dehydrated in ethanol (30%, 50%, 70%, 90%, and 2X 100%), and embedded in EPON resin before polymerization at 60 °C for 48 h. Ultrathin sections (~80 nm) were cut using a diamond knife and an Ultracut UC6 microtome (Leica Microsystems, Australia) and stained with uranyl acetate and lead citrate prior to imaging on a Hitachi HT7700 Transmission Electron Microscope at the Centre for Microscopy and Microanalysis (University of Queensland).

*Stable isotope incubation experiments*

To examine the effects of heat stress on inorganic carbon fixation by the sponge microbiome, sponge fragments from day 8 were incubated in artificial seawater prepared with $^{13}$C-labeled sodium bicarbonate (99% $NaH^{13}CO_3$) at a final concentration of 5 µM. Sponges were incubated in the light (80 µmol $m^{-2}$ $s^{-1}$) in sealed 1L chambers (800mL seawater with 200mL headspace) that were stirred to provide water flow and partially submerged in a flow-through water bath to maintain experimental temperatures (n=3 replicates per treatment). After 6h incubation, sponge fragments were transferred back to flow-through seawater for 10 min to remove any unincorporated $^{13}$C-labelled bicarbonate, rinsed in FSW, and sampled as described for FISH. Untreated sponges were sampled prior to the start of the experiment to determine background natural abundance values for $^{13}$C atom percent.

*CARD-FISH - NanoSIMS of sponge tissue sections*

Samples from T = 8 days (one control and one treatment sample for the thick morphotype) were processed for CARD-FISH NanoSIMS. Cryosections (5 µm) were prepared and transferred onto Indium Tin Oxide (ITO, Sigma, Darmstadt, Germany) wafers at the Histology Facility of the Vienna Biocenter (Vienna, Austria). In brief, fixed sponge samples were washed for 1 h in 1x PBS, stored overnight in 30% sucrose at 4°C, and subsequently embedded in 50% sucrose:Tissue-Tek O.C.T (Optimal Cutting Temperature) compound embedding medium (Sakura, Torrance, USA) overnight at 4°C. The embedded sponge samples were transferred to isopentane, frozen in liquid nitrogen, and kept at -80°C until cryosections were prepared. Regions of interest on the ITO wafers were marked using a laser microdissection microscope (Leica LMD 7000, Wetzlar, Germany). Prior to that, ITO wafers were submerged for 3 min each in 50%, 80%, and 96% ethanol to remove cryo-medium residue.

CARD-FISH was performed following Pernthaler et al. [6] with modifications described by Glasl et al. [7] using probes Arch915 [8] and AlfD729 with helper probes [7, 9], as well as negative controls using probe NON338 [10]. In brief, samples were permeabilised using lysozyme (10 mg $ml^{-1}$ in 0.05 mol $L^{-1}$ EDTA, 0.1 mol $L^{-1}$ Tris*HCl, pH 8; 60 min at 37°C), washed two times in deionized water, permeabilised using HCl (0.2 mol $L^{-1}$, 10 min, room temperature), and then again washed two times in deionized water, and subsequently dried. Endogenous peroxidases were inactivated using a freshly prepared 0.3% $H_2O_2$ solution in 1x PBS (30 min, room temperature) and samples were subsequently washed in deionized water and dried. Two consecutive rounds of hybridisation (each 3 h, 46°C, at the respective optimal formamide concentration, see Glasl et al. 2025 [7]) and signal amplification (each 30 min, 46°C; using Alexa488 and Alexa594 tyramides) were performed. Between hybridisations, the previously used probe was inactivated with 0.01 mol $L^{-1}$ HCl (10 min, at room temperature), followed by washing in deionized water twice. Samples were counterstained using DAPI (1 µg $ml^{-1}$, 10 min, room temperature), washed twice in deionized water, and air dried, before embedding in CitiFluor AF1 (EMS, Morgantown, USA). CARD-FISH and DAPI signals were visualised using a confocal laser scanning microscope (SP7, Leica, Wetzlar, Germany, equipped with a white light laser) across the entire volume of the sponge sections using z-stacks in combination with a maximum projection algorithm. After visualisation, the CitiFluor AF1 medium was removed by washing in deionized water three times for 5 min.

*NanoSIMS imaging and data analysis*

NanoSIMS measurements were performed on a NanoSIMS 50L instrument (Cameca, Gennevilliers, France) the NanoSIMS facility of the Centre for Microbiology and Environmental Systems Science, University of Vienna. Prior to data acquisition, analysis areas were pre-conditioned by rastering with a high-intensity, defocused $Cs^+$ beam following the extreme low ion impact energy (EXLIE) deposition procedure. Briefly, samples were first implanted at high energy (HE; 16 keV impact energy) using a 100 pA beam to a fluence of $5 \times 10^{14}$ ions $cm^{-2}$, followed by an EXLIE step at 50 eV with a 400 pA beam to a fluence of $5 \times 10^{16}$ ions $cm^{-2}$, and finished with a second HE step to an additional fluence of $2.5 \times 10^{14}$ ions $cm^{-2}$. The multicollection detector array was configured for simultaneous detection of the secondary ions $^{12}C^{12}C^-$, $^{12}C^{13}C^-$, $^{12}C^{14}N^-$, $^{12}C^{15}N^-$, $^{31}P^-$, and $^{32}S^-$. The instrument was tuned to a mass resolving power (MRP; Cameca definition) > 9,000 to ensure separation of $C_2^-$ and $CN^-$ species (slit combination: D1#4, ES#3, AS#2). Data were acquired as multilayer image stacks by sequential rastering of a finely focused $Cs^+$ primary beam (~1 pA), yielding an estimated lateral resolution of ~80 nm. Scanned areas were 40 × 40 to 45 × 45 µm², recorded at 512 × 512 pixels with a dwell time of 5 ms per pixel per cycle. For each area, 42–60 planes were collected. NanoSIMS images (n= 1 per treatment) were processed using the OpenMIMS plugin for ImageJ/Fiji (https://github.com/BWHCNI/OpenMIMS). Secondary ion intensities were corrected for dead time and quasi-simultaneous arrival effects, using sensitivity correction factors of 1.06 for $C_2^-$, 1.05 for $CN^-$, and 1.10 for all other ion species. Maps of $^{13}C/^{12}C$ enrichment were generated by taking the ratio of the drift corrected and stacked $^{13}C^{12}C^-$ and $^{12}C2^-$ images and displayed as hue-saturation-intensity (HSI) images. Individual microbial cells were identified based on $^{31}P^-$ secondary ion maps and correlated with CARD-FISH and DAPI signals to distinguish the three dominant symbionts. However, we were unable to align the $^{31}P^-$ images and CARD-FISH images for the heat stress sample. Instead, the ammonia-oxidizing-archaea were inferred from maps of $^{13}C$ enrichment as the only symbiont cells to assimilate inorganic carbon. Due to low numbers of the dominant archaea in the heat stress sample, single cell enrichments were not quantified and the difference in $^{13}C$ enrichment between treatments was compared qualitatively.

**Supplementary Note 1**

A total of 35,033 (thin) and 36,132 (thick) isogroups (i.e. genes) were included in the WGCNA analysis and were assigned to 17 (thin) and 16 (thick) modules. Each module consisted of genes that showed a similar expression pattern throughout the experiment (**Figure S3**). Module size varied from 7 to 5,190 genes for the thin and from 179 to 4,779 for the thick morphotype (**Figure S3A and C**). Each module was colour coded. For the thin morphotype, genes in the darkolivegreen (5,190 genes), darkslateblue (47), and orange (1,052) modules were positively correlated to temperature (i.e. increased expression under thermal stress), while genes in the blue (3,077) and orangered (284) modules were negatively correlated with temperature (**Figure S3A and C**). For the thick morphotype, genes in three modules were positively correlated with temperatures: ivory (324), lightcyan (2,996), and magenta (2,401, **Figure S3B and D**). Modules containing genes that were downregulated in response to heat stress were blue (4,779), plum2 (836), turquoise (3,846), and mediumpurple3 (686). As expected, select modules correlated to genotype rather than temperature (**Figure S3A and B**) and genes in these modules were omitted from further analyses. The darkslateblue module (thin morphotype) further correlated to time point and was also excluded from further analyses (**Figure S3A**).

**Supplementary Table descriptions**

**Table S1.** Sampling details of the study.

**Table S2.** Host reference transcriptome statistics and completeness for the thin and thick *I. basta* morphotype.

**Table S3.** Count data per *I. basta* sample after mapping of Tag-seq reads to the host reference transcriptomes.

**Table S4.** Differentially abundant genes per morphotype over time, including the PFAM annotation of the isogroup.

**Table S5.** Genes that are part of each enriched Gene Ontology term described in the manuscript, separated per individual and module (see **Figure S3** for the module-trait correlations).

**Table S6.** Relative abundance and taxonomy of identified amplicon sequence variants across samples.

**Table S7.** Amplicon sequence variants unique to the treatment samples (32 °C) at T = 8 days.

**Table S8.** Expressed proteins in the three thin and three thick *Ianthella basta* individuals. Each protein sequence was given a unique sequence ID (seqID). Information on the source assembly, whether the protein was captured in a MAG published in Engelberts et al. [27], and the function of the sequence are also included in the table.

### Supplementary Figures

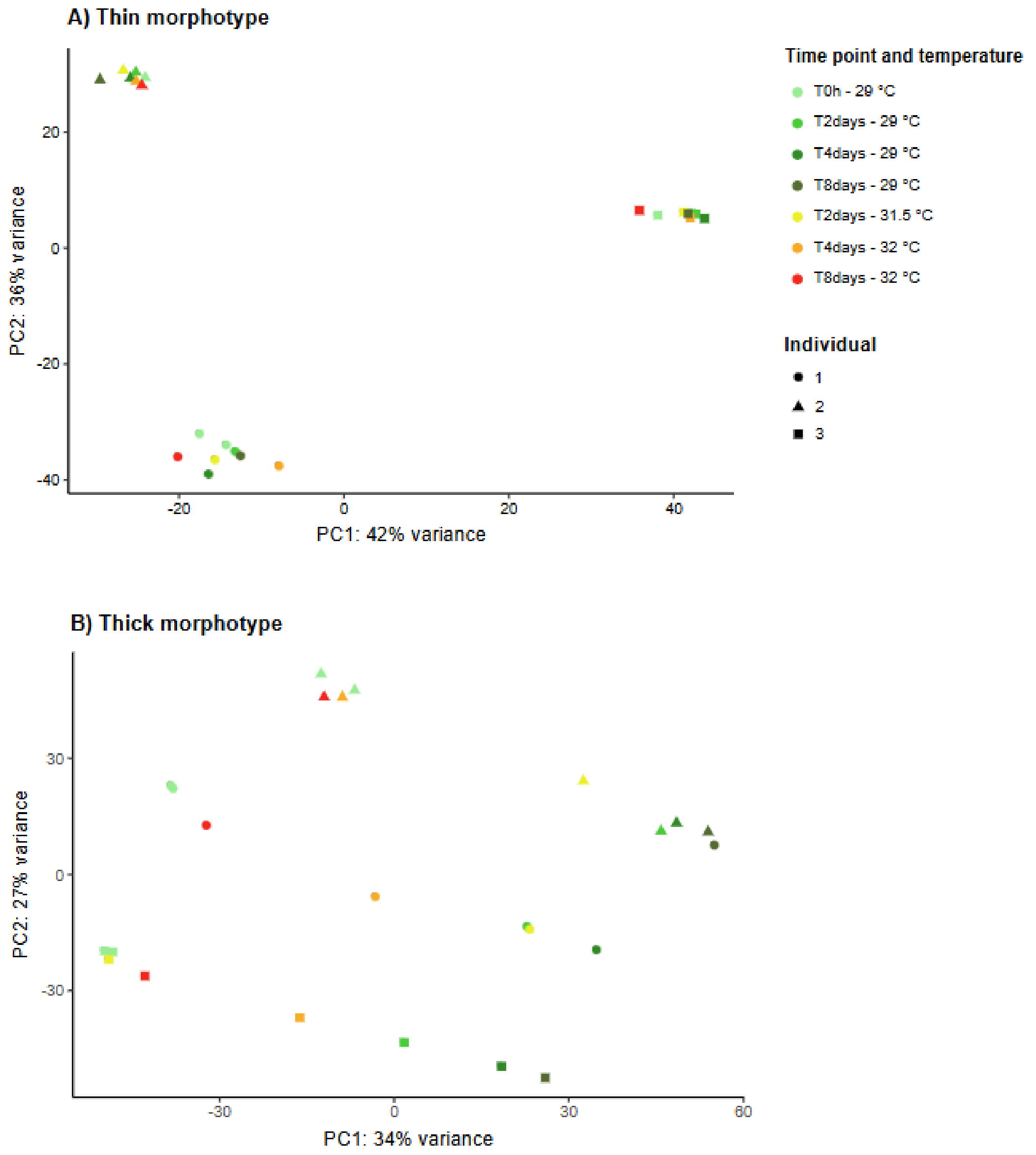


**Figure S1.** Principal component analysis (PCA) of the overall expression profile of *I. basta* individuals with a thin (A) and thick (B) morphotype.

A)

| Timepoint | Upregulated | | Downregulated | |
|---|---|---|---|---|
| | Thin | Thick | Thin | Thick |
| T = 0 h | 2 | 0 | 0 | 1 |
| T = 2 days | 17 | 43 | 0 | 0 |
| T = 4 days | 71 | 837 | 169 | 1520 |
| T = 8 days | 1670 | 2129 | 2114 | 3315 |

B) Thin morphotype

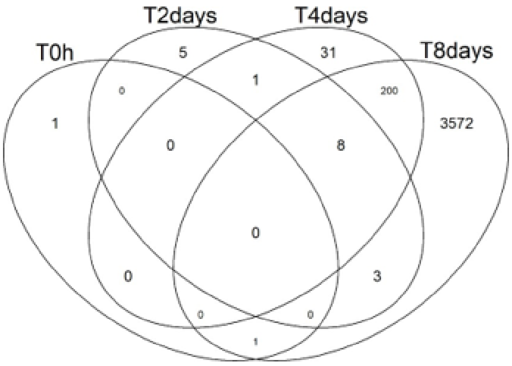


C) Thick morphotype

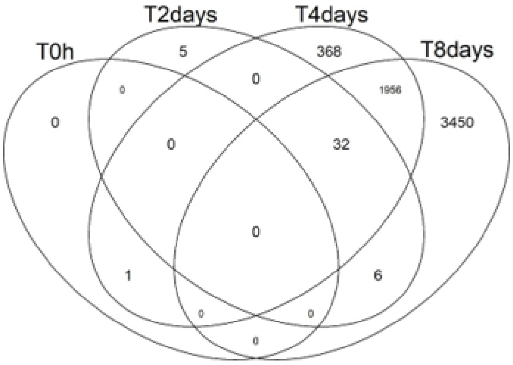


**Figure S2.** A) Number of significantly up- and downregulated genes in treatment samples versus control samples per time point. B-C) Venn diagram showing the overlap in differentially abundant genes per time point for the thin morphotype (B) and the thick morphotype (C). For instance, 239 of the 208 genes that were differentially expressed at T = 4 days in the thin morphotype remained differentially expressed at T = 8 days.

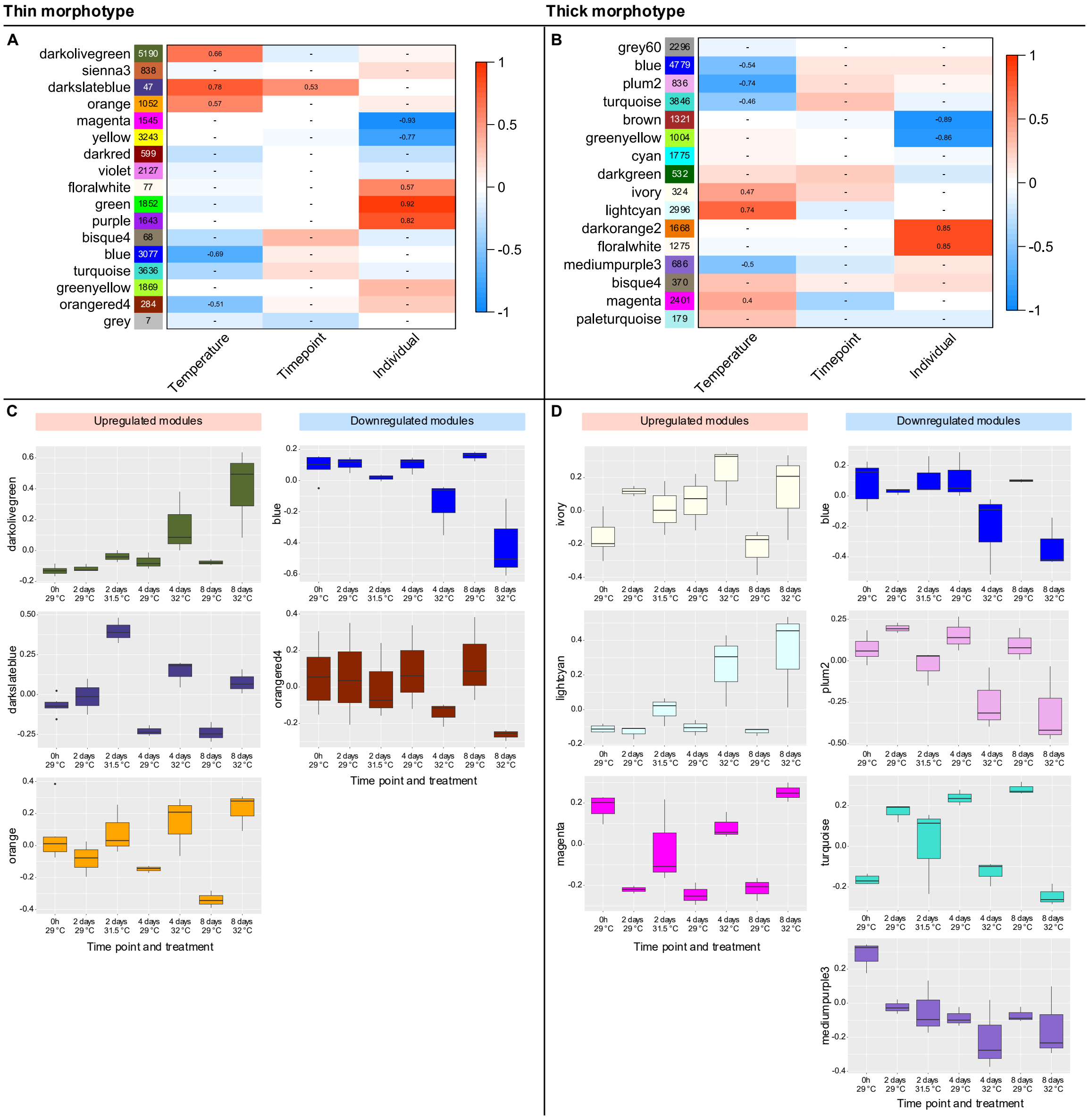


**Figure S3.** Heatmap of module-trait correlations for the thin (A) and thick (B) *I. basta* morphotype. Each row corresponds to a module eigengene. The module colour name is followed by the number of genes that are in the module. Red indicates a positive correlation and blue a negative correlation. Only significant correlation values are displayed. Module eigengene expressions are shown in individual boxplots below the heatmap (C and D), only including modules that are significantly correlated to temperature. Details on the Gene Ontology (GO) terms and genes that were significantly up- or downregulated in both morphotypes based on these modules can be found in **Table S4**.

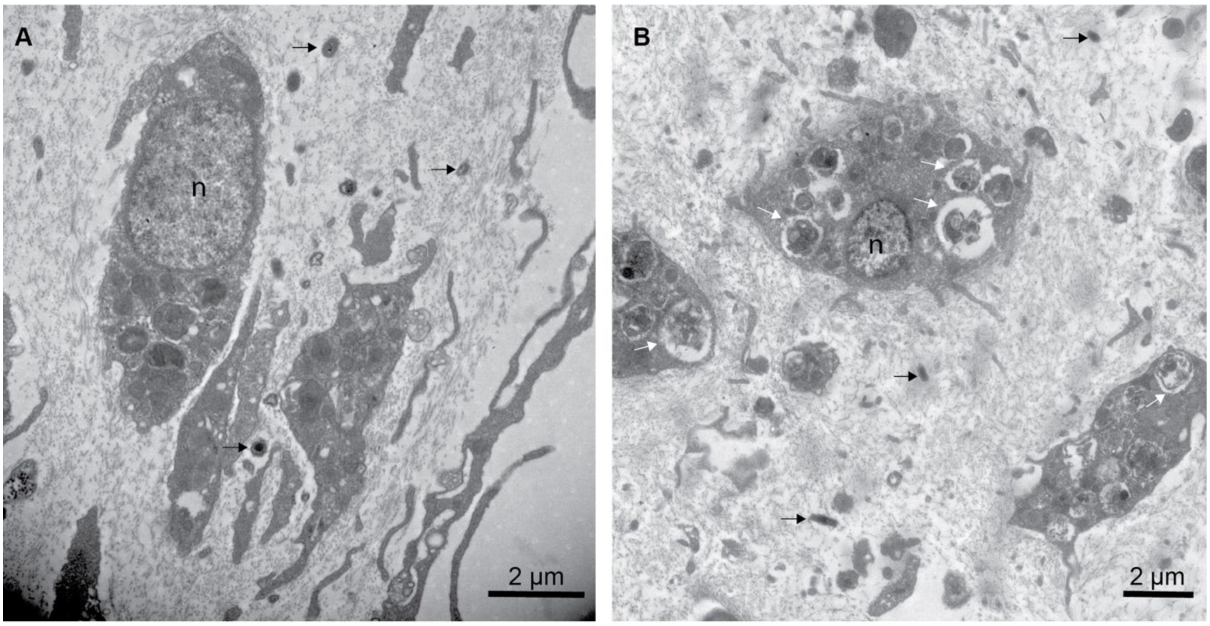


**Figure S4.** Transmission electron micrographs depicting *I. basta* tissue (thin morphotype) from the A) control and B) heat stressed treatments at day 8. The host sponge cells in the heat stressed contain numerous vacuoles of phagocytosed material indicative of late-stage apoptosis. Sponge nuclei (n), vacuoles containing phagocytosed material (white arrows), microbial cells (black arrows).

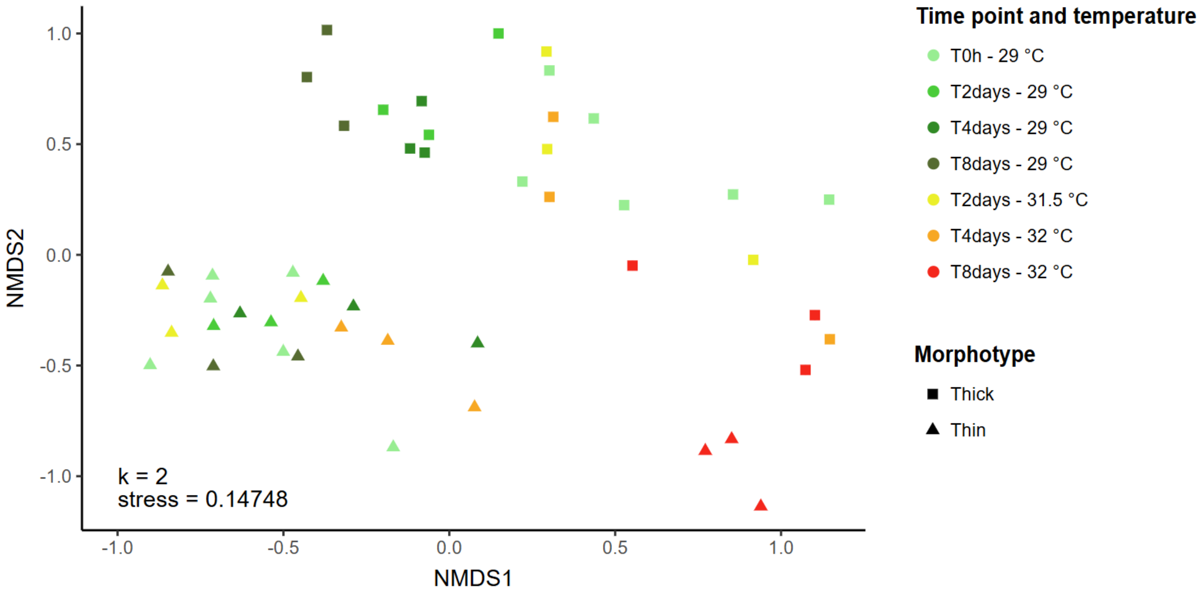


**Figure S5.** Non-metric multidimensional scaling plot displaying the (dis)similarity in the microbial community of *I. basta* individuals. The microbial communities of the thin and thick morphotypes vary in taxonomic composition and become more similar after 8 days of heat stress.

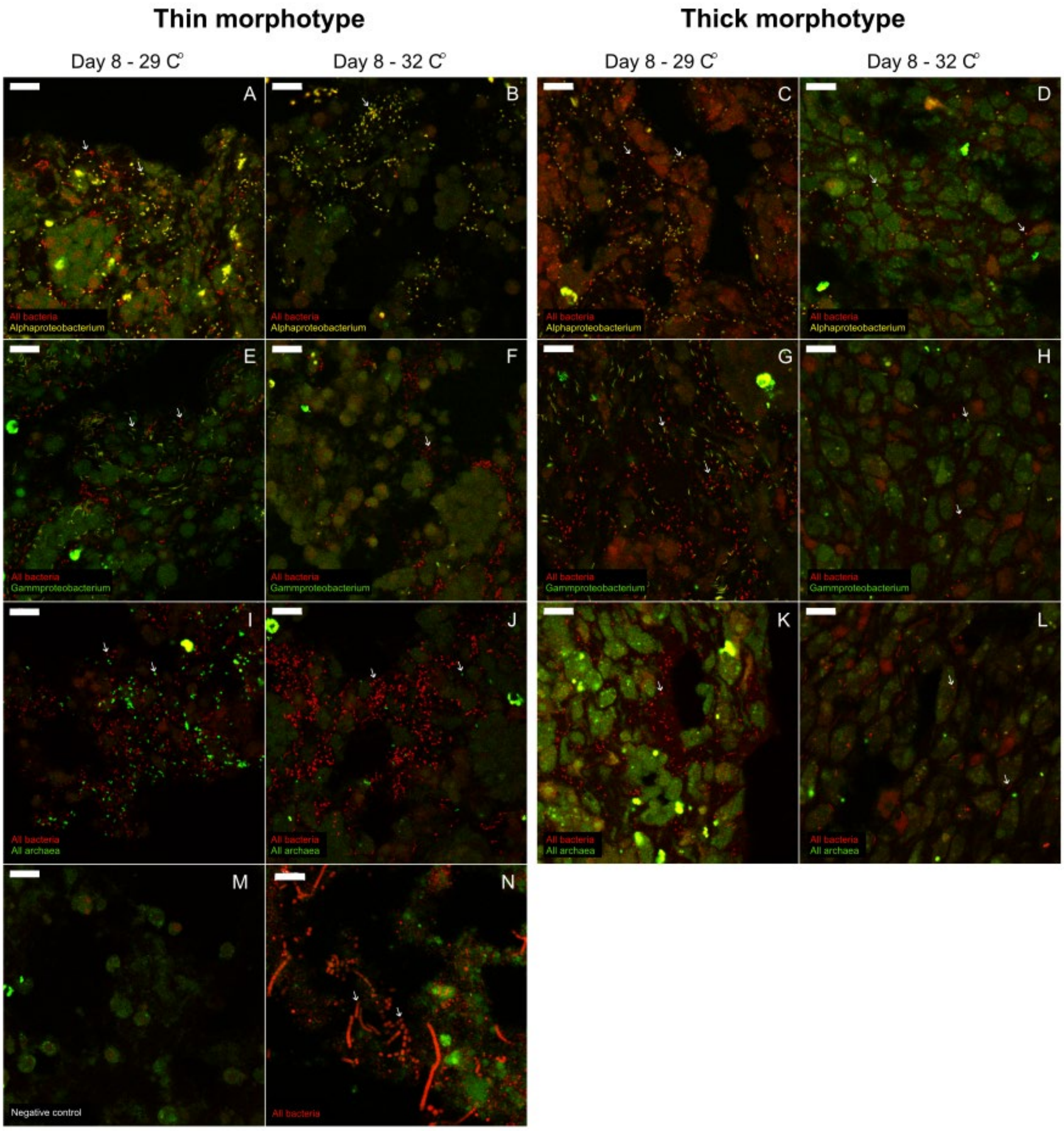


**Figure S6.** Fluorescence *in situ* hybridisation of a thin and thick morphotype of *I. basta* from either the control (A, C, E, G, I, K, M) or heat stressed treatment (B, D, F, H, J, L, N). Tissue was hybridised with A-D) the general EUBI-III probe in Cy3 (red) and the Alphaproteobacterium specific probe in Cy5 (yellow), E-H) the general EUBI-III probe in Cy3 (red) and the Gammaproteobacterium specific probe in Cy5 (yellow), I-L, N) the general EUBI-III probe in Cy3 (red) and the general ARC915 probe in Cy5 (green). M) Signals observed with control probe NON338 in Cy3 (red) and Cy5 (blue). Panel N shows the introduction of new microbial morphotypes after 8 days of heat stress. White arrows point towards bacteria and archaea. Scale bars are 10 μm.

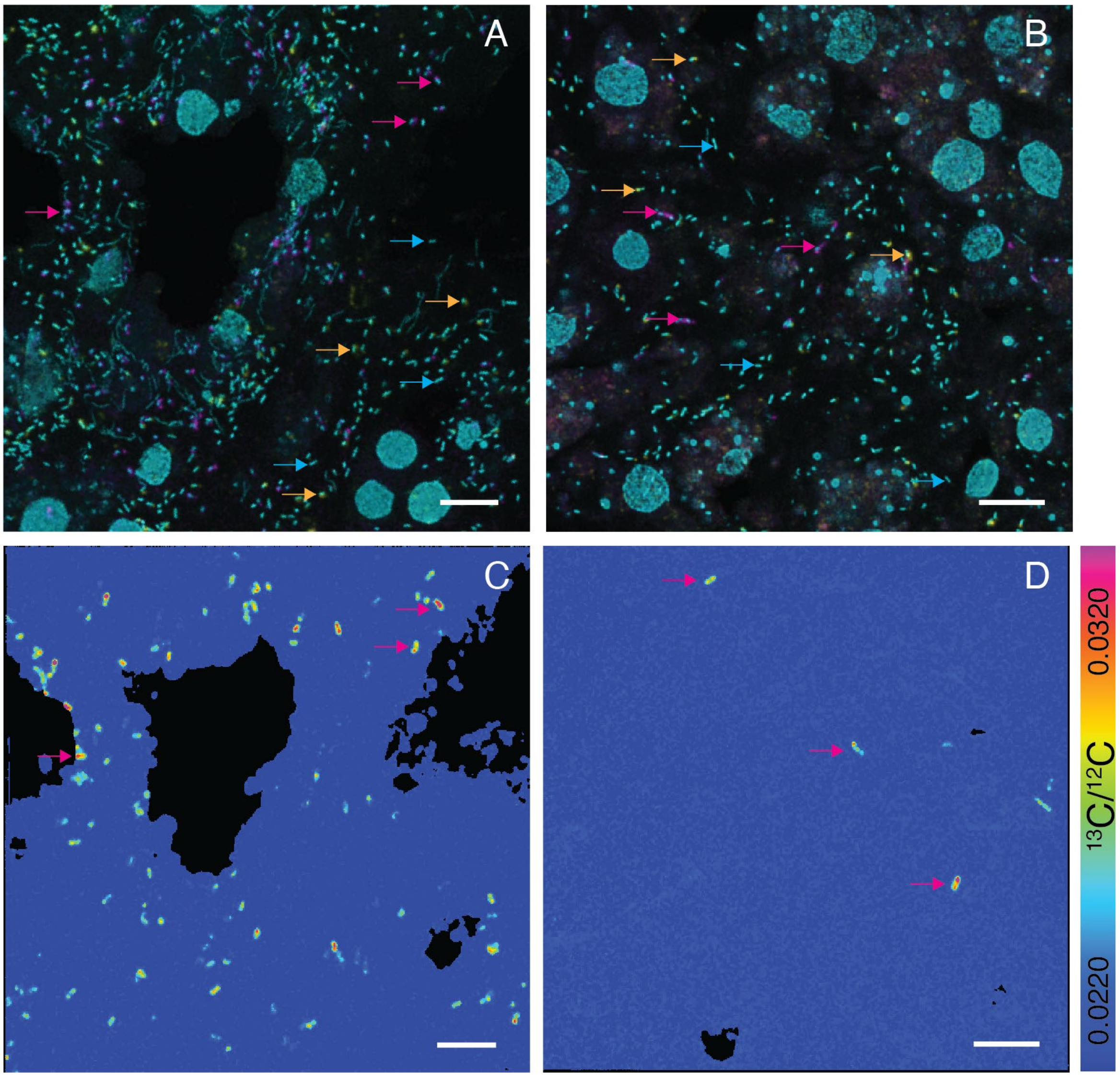


**Figure S7.** Single-cell enrichment of $^{13}C$ in the thick morphotype of *Ianthella basta* in the control and heat stress treatment after incubation with $^{13}C$-bicarbonate ($H^{13}CO_3^-$). CARD-FISH images of A) control and B) heat stress tissue hybridised with the archaeal probe ARC915 (magenta), the alphaproteobacterial probe AlfD729 (yellow), and counterstained with DAPI to show the sponge nuclei and rod-shaped gammaproteobacteria cells (cyan). Corresponding NanoSIMS hue-saturation-intensity images showing the distribution of $^{13}C/^{12}C$ enrichment in C) control and D) heat stress treatments. The colour scale represents enrichment relative to natural abundance ratios (blue). Scale bars are 5 µm.

### References


1. Grabherr, M.G., et al., *Trinity: reconstructing a full-length transcriptome without a genome from RNA-Seq data.* Nat Biotechnol, 2011. **29**(7): p. 644.
2. Simão, F.A., et al., *BUSCO: assessing genome assembly and annotation completeness with single-copy orthologs.* Bioinformatics, 2015. **31**(19): p. 3210-3212.
3. Kopylova, E., L. Noé, and H. Touzet, *SortMeRNA: fast and accurate filtering of ribosomal RNAs in metatranscriptomic data.* Bioinform, 2012. **28**(24): p. 3211-3217.
4. Eddy, S.R., *Accelerated profile HMM searches.* PLoS computational biology, 2011. **7**(10): p. e1002195.
5. R Core Team, *R: A language and environment for statistical computing.* 2013.
6. Pernthaler, A., J. Pernthaler, and R. Amann, *Sensitive multi-color fluorescence in situ hybridization for the identification of environmental microorganisms.* 2004.
7. Glasl, B., et al., *Branched-chain amino acid assimilation enables mixotrophy of ammonia-oxidizing archaeal sponge symbionts.* Sci Adv, 2026. **12**(36): p. eaef9450.
8. Stahl, D. and R. Amann, *Development and application of nucleic acid probes In: Stackebrandt E, Goodfellow M, editors. Nucleic acid techniques in bacterial systematics*. 1991, New York: John Wiley & Sons.
9. Moeller, F.U., et al., *Taurine as a key intermediate for host-symbiont interaction in the tropical sponge Ianthella basta.* ISME J, 2023: p. 1-16.
10. Wallner, G., R. Amann, and W. Beisker, *Optimizing fluorescent in situ hybridization with rRNA-targeted oligonucleotide probes for flow cytometric identification of microorganisms.* Cytometry, 1993. **14**(2): p. 136-143.
